# PolyCut: a computer programme for assignments of MS^*n*^ signals of metallo-biomolecules by considering the entire isotope patterns

**DOI:** 10.1101/089706

**Authors:** Zhenyu Shi, Chak Ming Sze

**Affiliations:** School of Chemistry and Bio21 Institute, University of Melbourne, Victoria 3010, Australia

**Keywords:** cisplatin, copper resistance protein C, antioxidant 1, copper chaperone proteins, tandem mass spectrometry

## Abstract

A computer programme ‘PolyCut’ has been written to assist mass spectrometry-based bioinorganic studies via top-down and bottom-up approaches in positive and negative ion modes. PolyCut is designed to cope with metalloproteins, metallo-polymers and biomolecules with heavy element(s) as one (or more) of the charging agents at defined but different oxidation states. It does automatic assignment by considering the entire experimental and simulated isotopic patterns of an ion rather than reading only the mono-isotopic and average *m/z* values (and/or mass values). The calculation of chemical formula and the simulation of isotopic pattern for an ion at various charge states require PolyCut to be informed of the user-defined elements, the user-defined abundance of isotopes for each element, the user-defined oxidation state of an element, the user-defined residues and sequence of a polymer, and the user-defined fragmentation (or digestion). The programme successfully assisted MS*^n^* studies of two copper proteins coordinated to platinum(II) centres (a functional group possessing an intrinsic positive charge rather than a neutral organic modification), published as *J*. *Biol*. *Inorg*. *Chem*. **2009**, *14(2)*, 163-165; *Metallomics* **2013**, *5(8)*, 946-954; and [more]. Besides polypeptides, polynucleotides and polysaccharides, PolyCut can be extended to other polymers using user-defined monomer residues. For the metallo-community.

Abbreviations

cisplatin: *cis*-Pt(NH_3_)_2_Cl_2_
en: 1,2-diaminoethane
bpy: 2,2’-bipyridine
equiv: equivalent
KPi: potassium phosphate
NH_4_OAc: ammonium acetate
ESI: electrospray ionisation
TOF: time-of-flight
FT-ICR: Fourier transformation cyclotron resonance
MS*^n^*: tandem mass spectrometry
CID: collision-induced dissociation
ECD: electron-captured dissociation

## Introduction

The latest TOF, FT-ICR and Orbitrap mass spectrometers feature extremely fast acquisition rates, combined with both high accuracy and resolution. Complex biomolecules often generate a large number of well-resolved isotope patterns derived from multiple fragmentation or digestion sites. Consequently, fast acquisition of data means that tedious and time-consuming visual analysis of their (tandem) mass spectra constitutes the rate-determining step in MS^n^-based studies of metalloproteins.

Visual analysis requires a high level of expertise [*Anal*. *Chem*. **2004**, *76(8)*, 2220-2230.]. However, a rich variety of MS software (free, commercial or provided on online workstations) is currently available for automatic analysis. A listing is given in Appendix 3.1.

Close examination of the listed softwares reveals that all programmes utilise the lightest isotope of a given element, disregarding all other heavier isotopes. In addition, the so-called ‘mono-isotopic’ species generated in this way from an isotope pattern is searched in an experimental (tandem) mass spectrum, ignoring all possible hetero-isotopic species. The user is responsible for ensuring that the ‘mono-isotopic’ species in the experimental isotope patterns is present at reasonably high relative abundance. The *m/z* spectrum will be ‘de-isotoped’ (figure 3.1) to have isotope peaks removed and converted to a numerical list called a ‘mass list’ of *m/z* values or masses of all mono-isotopic species so that an amino acid residue, a peptide sequence or a fragment is linked to a single datum only [*Proteomics* **2003**, *3(8)*, 1597-1610; *Anal*. *Chem*. **2004**, *76(8)*, 2220-2230.]. Information on the abundances (i.e., the Y-axis parameter in the *m/z* spectrum) is lost. [More]. The softwares listed in Appendix 3.1 perform analysis on a mass list but not on an entire *m/z* spectrum. Their application is effectively limited to bottom-up analysis or small peptides containing common elements such as C, H, N, O, S and P featuring mono-isotopes of reasonably high relative abundances. They still make many peptide mis-assignments and therefore require subsequent time-consuming visual analysis [*OMICS* **2002**, *6(2)*, 207-212.].

However, the hetero-isotopic patterns and their relative abundances provide important information and should be considered. The importance of the consideration of relative isotopic abundance of isotope peaks within an isotope pattern was stated recently [*Anal*. *Chem*. **2011**, *83(10)*, 3737-3743.]. The listed softwares cannot correctly assign signals arising from ions containing heavy elements of biological or medicinal relevance. These include Cr, Mo, W, Fe, Ru, Os, Pd, Pt, Cd, Hg, Pb, Se and Gd, all of which feature lightest isotope(s) of low abundance(s). Due to their low abundance, ions containing these isotopes may not be detected experimentally. The simple isotopic patterns of molecular ions become complicated when they are combined with heavy elements [*Metallomics* **2011**, *3(6)*, 550-565.].

Nevertheless, some of the softwares listed in Appendix 3.1 have been applied in studies of interactions of transition metal complexes of platinum and ruthenium with individual known proteins or with the contents of cells containing groups of unidentified proteins [*Chem*. *Commun*. **2001**, *(24)*, 2708-2709; *J*. *Biol*. *Inorg*. *Chem*. **2007**, *12(6)*, 883-894; *J*. *Anal*. *At*. *Spectrom*. **2008**, *23(3)*, 378-384; *J*. *Biol*. *Inorg*. *Chem*. **2008**, *13(3)*, 421-434; *ChemMedChem* **2008**, *3(11)*, 1696-1707; *Proteomics* **2011**, *11(21)*, 4174-4188.]. Their application to such systems can be summarized as follows:

1. One of the highly abundant isotopes of the metal element is defined manually to be the ‘lightest isotope’ (e.g. ^190^Pt, ^192^Pt, ^194^Pt, **^195^Pt**, ^196^Pt, ^198^Pt; ^96^Ru, ^98^Ru, ^99^Ru, ^100^Ru, ^101^Ru, **102Ru**, ^104^Ru. Post Pt & Ru isotope patterns.).
2. Mass units or hydrogen atoms are substracted manually in order to adjust for the anticipated metal oxidation states (e.g. by entering 'Ru − 2H' instead of Ru^2+^ where Ru means Ru^0^).
3. Re-examination, by visual analysis, of the interpretations obtained from the softwares.

The advantages and deficiencies of the programmes Mascot and SEQUEST are considered here as they have been applied widely on analysis of metallo-biomolecular systems (Refs: choose only the most important from the following refs for software on analysis of metallo-biomolecular systems:

[*Rapid Commun*. *Mass Spectrom*. **2002**, *16(10)*, 933-935; *ChemBioChem* **2005**, *6(10)*, 1788-1795; more.]

[*Chem*. *Commun*. **2001**, *(24)*, 2708-2709.***]

[*J*. *Anal*. *At*. *Spectrom*. **2008**, *23(3)*, 378-384. In this paper, Mascot was applied to sequence *apo*-protein only. Assignment of Pt-containing signals totally via visual analysis.]

[*J*. *Am*. *Soc*. *Mass Spectrom*. **2007**, *18(7)*, 1324-1331.]

[More?]

**Cite the following papers for software analysis on transition metal / metallo-biomolecular studies (anymore?)**:

*J*. *Biol*. *Inorg*. *Chem*. **2007**, *12(6)*, 883-894.

*J*. *Biol*. *Inorg*. *Chem*. **2008**, *13(3)*, 421-434.

*ChemMedChem* **2008**, *3(11)*, 1696-1707.

[More?]

Programme SEQUEST has been applied to identify unknown proteins with bound transition metal ions [*J*. *Biol*. *Inorg*. *Chem*. **2007**, *12(6)*, 883-894; *J*. *Biol*. *Inorg*. *Chem*. **2008**, *13(3)*, 421-434.]. The metallated sequences were searched and identified automatically according to the metallo-modifications (only the most abundant isotope for each element were considered and it was necessary to manually compensate the 2+ charge of Ru^2+^ / Pt^2+^ and the 1− charge of an inorganic anionic ligand) on residues defined by the user. The authors in turn visually compared the experimental isotope patterns against simulations for the metallated sequences found by SEQUEST to confirm the results. Did this paper only looked at backbone *b*/*y’* fragmentations?

The tandem mass spectra in [*J*. *Biol*. *Inorg*. *Chem*. **2008**, *13(3)*, 421-434.] are in the form of ‘Data Dependent Acquisitions’ containing MS^1^ master scans. Only single scan MS/MS were recorded for each precursor ion. No averaging is possible and the experimental isotope patterns deviate significantly from simulations. This aspect may be made good by using the linear ion trap of the Finnigan LTQ to collect several MS/MS scans per second. In [*J*. *Biol*. *Inorg*. *Chem*. **2008**, *13(3)*, 421-434.], NH_3_ losses observed in platinated sequence?

The programmes Mascot and SEQUEST were also in [*ChemMedChem* **2008**, *3(11)*, 1696-1707.]. The approach is described below:

a. The software does not generate theoretical isotope patterns and does not consider experimental isotope patterns. No comparisons between the theoretical and experimental isotope patterns are possible. The user must generate theoretical isotope patterns manually and compare those with experimental patterns that are claimed to contain metal ion(s) by the software based on searches for ‘mono-isotopic’ species only. The user must also use his/her judgement to eliminate false positives or negatives. Mascot and SEQUEST were designed primarily for sequencing of peptide fragments. However, not all MS/MS spectra contain the necessary sequential information but they enable identifications of metal ion binding sites. In addition, the software cannot identify ions resulted from unusual side-chain fragmentations.
b. The software assumes that hydrogen is the only element capable of losing electrons and that all metal elements are neutral. Such manual adjustment is also popular in other MS softwares including the software packages MassHunter Qualitative Analyses (Agilent, Version B.0301 Build 3.1.346.14) and Xcalibur (Thermo Electron, version 2.0 SR1) which are able to generate theoretical isotope patterns according to user-entered molecular formula of each MS or MS*^n^* signal.
c. The software assumes that protons are the only source of positive charges in spectra generated by electrospray ionisations. Users need to manually subtract proton(s) from ions containing fixed positive charge(s) from other elements. Some programmes cannot analyse mass spectra in the negative-ion-mode.

This paper introduces a new programme called ‘PolyCut’ to overcome the difficulties discussed above.

## Experimental

### Materials

Chemicals, enzymers, solvents, etc.

### Samples

Define denaturing and non-denaturing conditions. Protease-digested samples.

### Mass spectrometry

Machines, ESI-TOF & FTICR experiments.

### Analytical basis of the programme ‘PolyCut’

PolyCut is coded using Microsoft^®^ Visual Basic 2010. A complete set of files for normal functioning of PolyCut contains individual files <u>PolyCut.exe</u>, <u>PolyCut.vshost.exe</u>, <u>PolyCut.vshost.exe.manifest</u>, <u>Element.XML</u>, <u>ElmAccl</u> <u>.XML</u>, <u>Molecule.XML</u>, <u>PolyCut.XML</u> and user-generated <u>.splt</u>. Their functions are explained in the following sections. Calculation of isotopic abundance distributions can be referred to [*Methods Enzymol*. **1990**, *193*, 882-886.]. Users are required to define a variety of primary settings:

1. *The atomic mass (referenced to ^12^C) and relative abundance of each isotope of any element present*. <u>Element.XML</u> defines these values which may be changed and saved from day to day if required, e.g., for isotopically labelled systems. Updates can be obtained via WebElements (http://www.webelements.com/index.html). A portion of <u>Element.XML</u> is shown in figure 3.2. The current <u>Element.XML</u> file contains the atomic masses and relative abundances of each isotope of the elements listed in table 3.1. These are the values applied throughout the analysis presented in X.
2. *The symbols and the molecular formulae for the monomer residues in a polymer*. It may be applied to a polypeptide, polynucleotide, polysaccharides or any molecular ion that may be fragmented/digested by any means as soon as the isotope peaks in the detected isotope patterns are resolved below 50% of the isotope peak height (figure 3.3) (this is the definition of a peak: if above 50%, the entire isotope pattern is treated as a single peak). <u>Molecule.XML</u> defines the one-letter (capital) abbreviations and the chemical formulae for each monomer residue. The current <u>Molecule.XML</u> file is shown in figure 3.4 containing one-letter abbreviations and chemical formulae for the 20 standard amino acid residues which are applied throughout the analysis of X (add description of figure 3.5). Add: one atom label or label eth with a specific isotope of any element.
3. *The sequence of the polymer*. This is entered as shown in figure 3.6. The button ‘Peptide Format’ converts it to the PolyCut-readable form, as shown in figure 3.7.
4. *The ionisation mode in as ESI or MALDI*.
5. *The fragmentation/digestion site(s) on a polymer and the molecular structures of the resulting ions* using regular expressions (Visual Studio) even if the molecular fragmentation/digestion mechanisms are unknown or hypothetical and are not limited to the peptide backbone. For instance, the following peptide fragment ions can be efficiently recognised as arising from the given experiments: the *a*, *b* and *y’* peptide fragment ions from low-energy CID; the *d*, *v*, and *w* peptide fragment ions from high-energy CID; and the *a*•, *y’*, *c’*, *z*• peptide fragment ions from ECD can be efficiently recognised. This feature requires users to have a high level of expertise in fragmenation behaviours of biopolymers in the gaseous phase. The relevant portion of the user-interface of PolyCut for this feature is shown in figure 3.8:

i. Select a command for each row in the Command column. <u>Command ‘cidfrag’</u>: to make one cut only in the single sequence within a pair of adjacent residues specified in the Pattern column. <u>Command ‘split’</u>: to make multiple cuts in the single sequence within pairs of adjacent residues specified in the Pattern column, with missing cuts not being allowed. <u>Command ‘digest’</u>: same as Command ‘split’, but with missing cuts allowed. <u>Command ‘attach’</u>: to add a functional group to specific residue(s) specified in the Pattern column. <u>Command ‘bridge’</u>: bridged by a functional group; this command can be extended to branched polymers. Commands ‘split’, ‘digest’ and ‘cidfrag’ are defined to produce one cut(s) within a pair(s) of adjacent residues only. The formula of an amino acid residue is defined in the current <u>Molecule.XML</u> file as (NH)CR(CO) where R is a functional group. Hence, if one cut is made within a pair of glycine residues in a tetrapeptide XaaGlyGlyXaa, a N-terminal fragment with chemical formula Xaa(NH)CH_2_(CO) and a C-terminal fragment with chemical formula (NH)CH_2_(CO)Xaa are produced.
ii. In the Pattern column, specify the location of the cut(s) for the commands ‘split’, ‘digest’ and ‘cidfrag’ or specify the residue(s) with a functional group for the command ‘attach’ via regular expression. For details of regular expressions, visit http://msdn.microsoft.com/en-us/library/2k3te2cs(VS.80).aspx.
iii. Enter a chemical formula in the Front column to fine tune the formula of the N-terminal fragment to, for example, that of a peptide digested from a protein, the formula of the *b* fragment in a *b*/*y’* pair from low-energy CID, the formula of the *a*• fragment in a *a*•/*y’* pair or the formula of the *c’* fragment in a *c’*/*z*• pair from ECD. This column only works for the commands ‘split’, ‘digest’ and ‘cidfrag’.
iv. The Rear column functions the same as the Front column to fine tune the formula of the C-terminal fragment.
v. Enter a chemical formula in the Additional column to allow a functional group located on a specified residue(s). This column only works for the commands ‘attach’ and ‘bridge’.

Practical applications of the user-interface shown in figure 3.8 are discussed in detail on figures A3.2.1 to A3.2.16 in Appendix 3.2. They appear throughout the analysis in X.
6. *The modification of the polymer involving any elements* such as metallation or neutral losses.
7. *The valency of metal ions and the charges of any fixed-charge functional groups* since not all charges originate from gain/loss of protons in electrospray ionisation. Figure A3.2.4 in Appendix 3.2 is an example assigning a fixed charge to a modification.
8. *The highest possible charge-state for the theoretical chemical species generated and the criteria for an ‘assignment match’*. The latter involves comparisons of the error in the *m/z* and the error in the relative intensity of each detected isotope peak in an experimental isotope pattern relative to simulation. When the experimental and the simulated isotope pattern are overlaid, the relative intensity of the most intense detected isotope peak in the experimental isotope pattern and that of the corresponding isotope peak in the simulation are defined as 100%. The relevant portion of the user-interface of PolyCut for this feature is shown in figure 3.9.

In accordance with the above user-defined primary settings, PC simulates isotope patterns of all theoretical fragmented/digested ions at all user-defined charge states in the Centroid mode. It imports a .txt file containing data points on *m/z* (X-axis) and absolute intensity (Y-axis) exported for a (tandem) *m/z* spectrum acquired by any mass spectrometers and their acquisition softwares in Profile (Gaussian) mode. PolyCut defines the *m/z* value of a detected isotope peak as the *m/z* at the mid-width of the peak at half-height (figure X). The defined *m/z* may fall between the those of two adjacent exported data points that shape the peak. In this case PolyCut picks their average value as the *m/z* value of the isotope peak. The accuracy of the defined *m/z* value depends on the exported precision (i.e. the number of exported decimal places) and the exported frequency of the data points that shape the isotope peak.

## Results and discussion

### Capability of programme PolyCut

To make assignments, PolyCut firstly searches for the detection of the most abundant simulated isotope peak in the imported *m/z* spectrum, disregarding whether that peak is a mono-isotope or not. The peak is regarded as detected if its simulated *m/z* value falls within the *m/z* range covered by the width of an experimental isotope peak at its half height. Secondly, PolyCut overlays the entire experimental isotope pattern against the simulation and examines all other detected isotope peaks according to user-defined limits to the errors in the *m/z* values and in the relative intensities referenced to the simulation.

PolyCut expedites the analysis of a large amount of ions derived from physico-chemical fragmentations (such as CID and ECD) and chemical/biochemical cleavages (e.g. by trypsin, GluC and CNBr). PolyCut can analyse (tandem) *m/z* spectra generated from both top-down and bottom-up approaches in positive- or negative-ion modes for ions containing *any* heavy elements and *any* modifications with fixed-charges. However, it is designed to analyse experimental isotope patterns with isotope peak resolution at 50% peak height or below, no matter whether the signal is detected by high-resolution FTICR, Orbitrap and TOF spectrometers, or low-resolution Ion-Trap spectrometers. With isotope peaks resolved above 50% peak height, the entire experimental isotope pattern will be regarded as a single isotope peak and no analysis can be proceeded.

The potential of PolyCut is demonstrated in X (more) via its performance in making assignment matches for fragmented/digested ions. Its predictions will be compared to visual analyses (and also e.g. Mascot) with different examples. Give a figure of overlay as an example.

The programme:

a. searches for the most intense peak (not the mono-isotopic peak) within an isotope pattern;
b. searches for *m/z* values, not mass values;
c. considers the entire isotope pattern and does not perform deconvolution nor does it ‘de-isotope’ the patterns;
d. automatically generates a list of theoretical fragments/digested peptides at different charge states with or without modifications, calculates their chemical formulae and creates their theoretical isotope patterns which are overlaid against the experimental ones during analysis;
e. allows the user to view the list, the chemical formulae, the theoretical isotope patterns and the overlaps mentioned in (d) by single-clicks on buttons (more: entire *m/z* spectrum labelled with IDs, assignment hits labelled and arranged, the simulated and experimental *m/z* values for every isotope peaks, errors in ppm, absolute intensities);
f. is not limited to C, H, N, O, S and P, and can be applied to elements with lightest isotope(s) at very low abundance(s) (e.g., Cr, Mo, W, Fe, Ru, Os, Pd, Pt, Cd, Hg, Pb, Se and Gd);
g. can be applied to top-down *m/z* spectra.

Users are required to impose the following conditions:

a. The isotope abundance(s) of element(s) in their samples and so PolyCut can be applied to isotope-labelled samples.
b. The number(s) and type(s) of atom(s) on a monomer residue.
c. The polymer sequence (which could be absent in a database or a variant sequence with unusual amino acid residues).
d. The ionization methods (ESI, FAB, etc).
e. The ionization mode (positive- or negative-ion).
f. The fixed-charge(s) on given element(s) (e.g., transition metal ions in different oxidation states).
g. The digestion or fragmentation mechanisms, even if only hypothesised.

In addition, PolyCut:

a. distinguishes between ions *a* from CID and *a*• from ECD where both have different mono-isotopic *m/z* values in every charge state but are shown to be the same in ProteinProspector v 5.6.2 (http://prospector2.ucsf.edu).
b. can be applied to all polymers including polypeptides, polynucleotides and polysaccharides, including cyclic and branched polymers.
c. cannot be applied for sequencing (including *de novo* sequencing) and protein identification and will not search for a sequence from any database.

Mention the problem of ‘MS1 master scan’. PolyCut cannot adapt files created by Data Dependent Acqusition.

### ESI-TOF experiments

### ESI-FTICR experiments

## Conclusion

The new programme ‘PolyCut’ has been developed to overcome the difficulties discussed in the introduction section:

a. It simulates full isotope patterns of all theoretical ions according to user-defined protein sequence, charge states, and the site plus the mechanism of fragmentation or digestion.
b. It recognises the valency of transition metals on the ions.
c. It imports a .txt file containing parameters of data points on *m/z* (X-axis) and absolute intensity (Y-axis) exported for a (tandem) *m/z* spectrum acquired by any mass spectrometers.
d. It firstly searches for the detection of the most abundant simulated isotope peak in the imported *m/z* spectrum. If the peak is detected, it overlays the entire experimental isotope pattern against the simulation and examines all other detected isotope peaks according to user-defined allowance of the errors in *m/z* and relative intensity referenced to the simulation before claiming an assignment match.

It expedites the analysis of a large amount of ions derived from CID and ECD, and from trypsin and GluC digests, effectively applied to (tandem) *m/z* spectra generated from both top-down and bottom-up approaches for ions containing charged platinum(II) centre(s).

Now distinguish mechanism via chemical formula of precursor and product ions.

Need add distinguish mechamism via actual chemical structure of precursor and product ions. [More].

Also make one figure to explain how to analyse MS^3^ *m/z* spectrum.

Make PolyCut able to handle a sequence of only one residue isotopically labelled.

**Figure 1.**
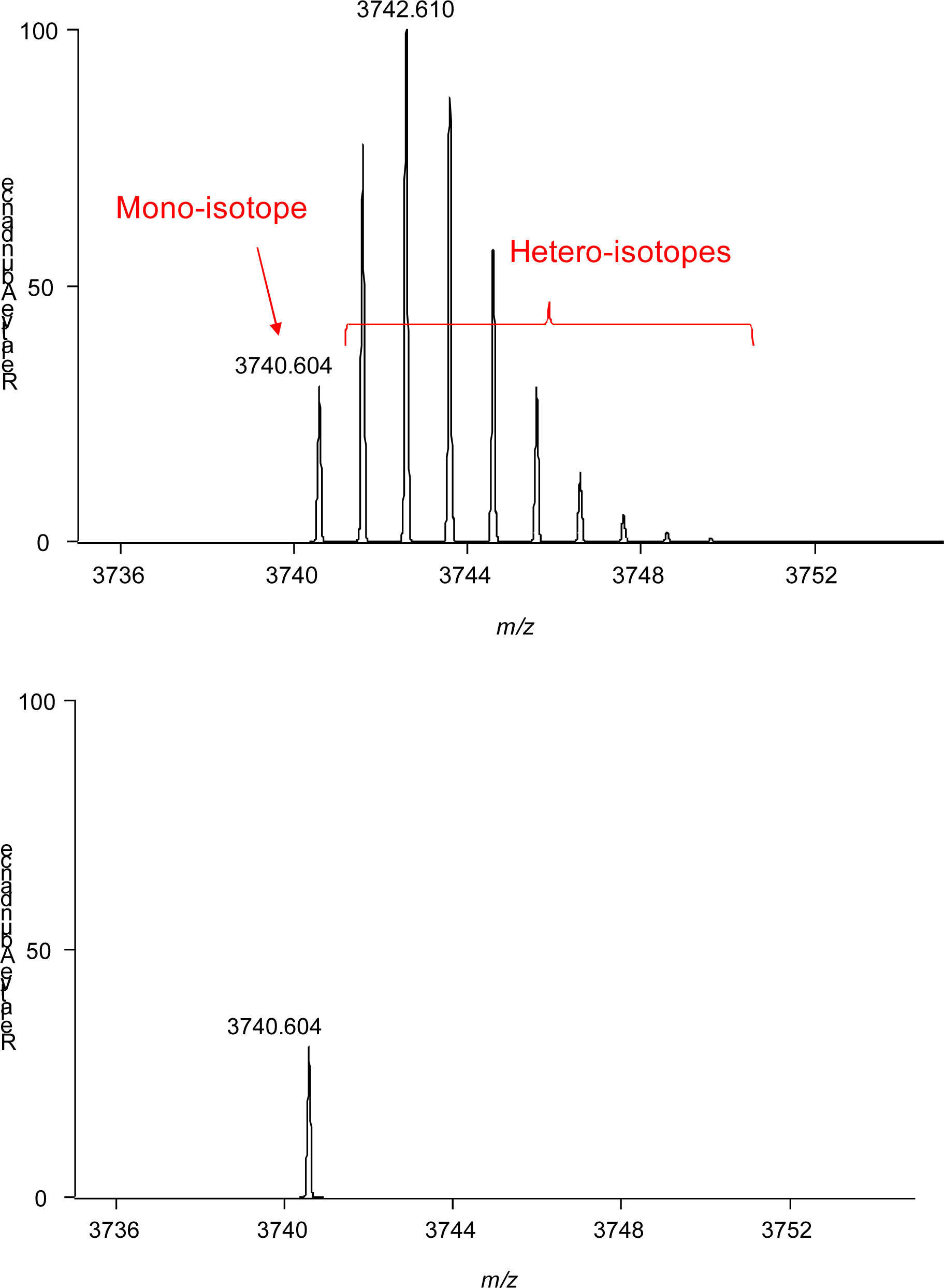
An isotope pattern (upper panel) and its de-isotoped form (lower panel) of a polypeptide containing 20 tryptophan residues at 1+.

**Figure 2.**
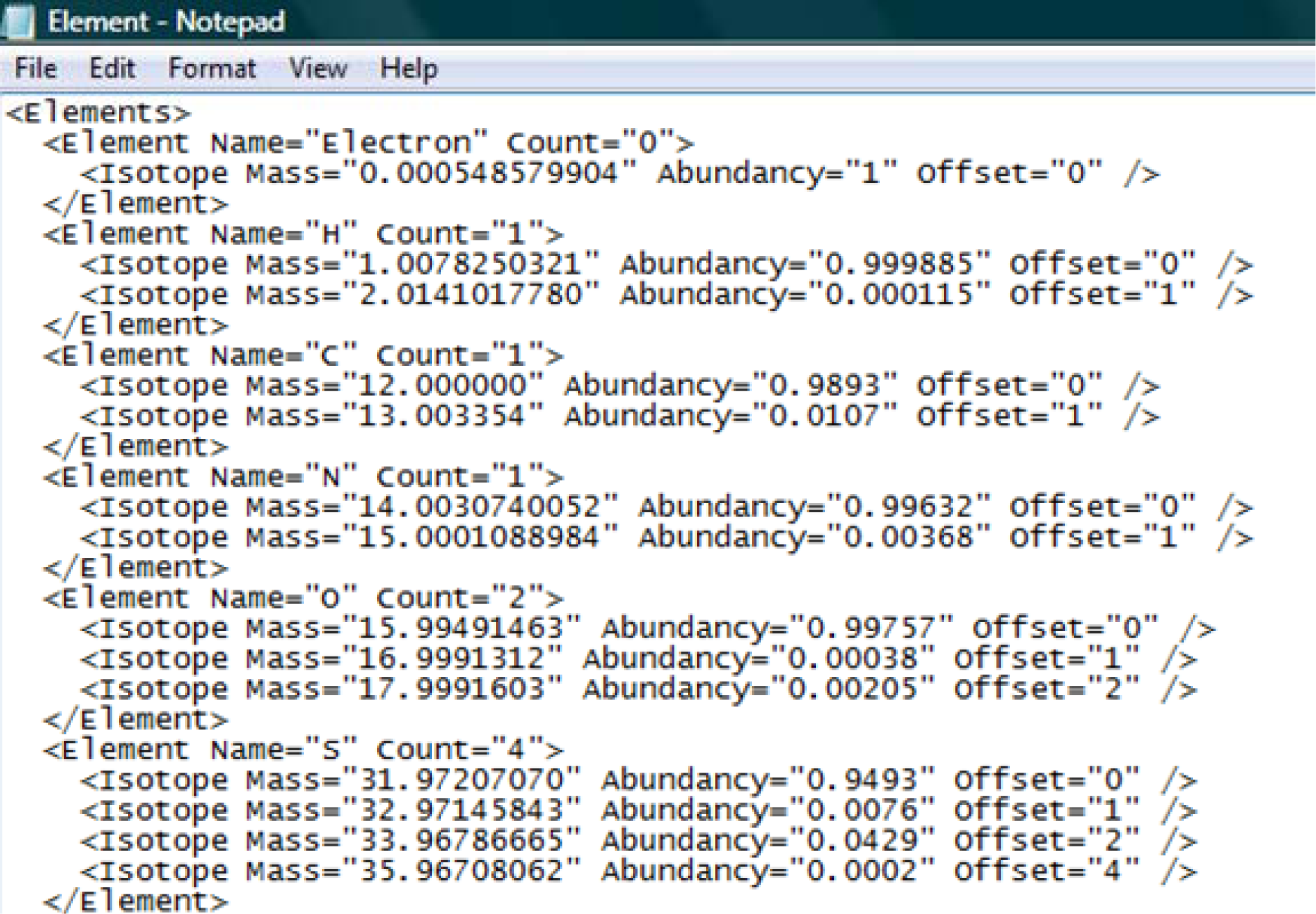
A portion of <u>Element.XML</u>.

**Table 1.** The relative masses and abundances of the electron and all natural isotopes of selected elements currently implemented in PolyCut. They are all stable isotopes except ^40^K. Non-natural isotopes, no matter whether they are stable or radioactive, are not involved. All data were obtained on X from http://en.wikipedia.org/wiki/Electron_rest_mass and http://www.webelements.com/index.html.

|  | relative rest mass | abundance<br>(%) |  |
| --- | --- | --- | --- |
| $e^-$ | 0.00054857990943 | 100 | |

| isotope | relative atomic mass<br>(amu) | abundance<br>(%) | nuclear spin |
| --- | --- | --- | --- |
| $^{12}\text{C}$ | 12.000000 | 98.93 | 0 |
| $^{13}\text{C}$ | 13.003354 | 1.07 | 1/2 |
| $^1\text{H}$ | 1.0078250321 | 99.9885 | 1/2 |
| $^2\text{H}$ | 2.0141017780 | 0.0115 | 1 |
| $^{14}\text{N}$ | 14.0030740052 | 99.632 | 1 |
| $^{15}\text{N}$ | 15.0001088984 | 0.368 | 1/2 |
| $^{16}\text{O}$ | 15.99491463 | 99.757 | 0 |
| $^{17}\text{O}$ | 16.9991312 | 0.038 | 5/2 |
| $^{18}\text{O}$ | 17.9991603 | 0.205 | 0 |
| $^{32}\text{S}$ | 31.97207070 | 94.93 | 0 |
| $^{33}\text{S}$ | 32.97145843 | 0.76 | 3/2 |
| $^{34}\text{S}$ | 33.96786665 | 4.29 | 0 |
| $^{36}\text{S}$ | 35.96708062 | 0.02 | 0 |
| $^{31}\text{P}$ | 30.9737620 | 100 | 1/2 |
| $^{35}\text{Cl}$ | 34.968852721 | 75.78 | 3/2 |
| $^{37}\text{Cl}$ | 36.96590262 | 24.22 | 3/2 |
| $^{39}\text{K}$ | 38.9637074 | 93.2581 | 3/2 |
| $^{40}\text{K}$ | 39.9639992 | 0.0117 | 4 |
| $^{41}\text{K}$ | 40.9618254 | 6.7302 | 3/2 |
| $^{23}\text{Na}$ | 22.9897677 | 100 | 3/2 |
| $^{190}\text{Pt}$ | 189.959917 | 0.014 | 0 |
| <sup>192</sup>Pt | 191.961019 | 0.782 | 0 |
| <sup>194</sup>Pt | 193.962655 | 32.967 | 0 |
| <sup>195</sup>Pt | 194.964766 | 33.832 | 1/2 |
| <sup>196</sup>Pt | 195.964926 | 25.242 | 0 |
| <sup>198</sup>Pt | 197.967869 | 7.163 | 0 |
| <sup>63</sup>Cu | 62.9295989 | 69.17 | 3/2 |
| <sup>65</sup>Cu | 64.9277929 | 30.83 | 3/2 |
| <sup>64</sup>Zn | 63.9291448 | 48.63 | 0 |
| <sup>66</sup>Zn | 65.9260347 | 27.90 | 0 |
| <sup>67</sup>Zn | 66.9271291 | 4.10 | 5/2 |
| <sup>68</sup>Zn | 67.9248459 | 18.75 | 0 |
| <sup>70</sup>Zn | 69.925325 | 0.62 | 0 |
| <sup>107</sup>Ag | 106.905092 | 51.839 | 1/2 |
| <sup>109</sup>Ag | 108.904756 | 48.161 | 1/2 |
| <sup>197</sup>Au | 196.966543 | 100 | 3/2 |
| <sup>106</sup>Cd | 105.906461 | 1.25 | 0 |
| <sup>108</sup>Cd | 107.904176 | 0.89 | 0 |
| <sup>110</sup>Cd | 109.903005 | 12.49 | 0 |
| <sup>111</sup>Cd | 110.904182 | 12.80 | 1/2 |
| <sup>112</sup>Cd | 111.902757 | 24.13 | 0 |
| <sup>113</sup>Cd | 112.904400 | 12.22 | 1/2 |
| <sup>114</sup>Cd | 113.903357 | 28.73 | 0 |
| <sup>116</sup>Cd | 115.904755 | 7.49 | 0 |
| <sup>196</sup>Hg | 195.965807 | 0.15 | 0 |
| <sup>198</sup>Hg | 197.966743 | 9.97 | 0 |
| <sup>199</sup>Hg | 198.968254 | 16.87 | 1/2 |
| <sup>200</sup>Hg | 199.968300 | 23.10 | 0 |
| <sup>201</sup>Hg | 200.970277 | 13.18 | 3/2 |
| <sup>202</sup>Hg | 201.970617 | 29.86 | 0 |
| <sup>204</sup>Hg | 203.973467 | 6.87 | 0 |

**Figure 3**. Definition of the height and the mid-width at half height of an isotope peak within an isotope pattern.

**Figure 4.**
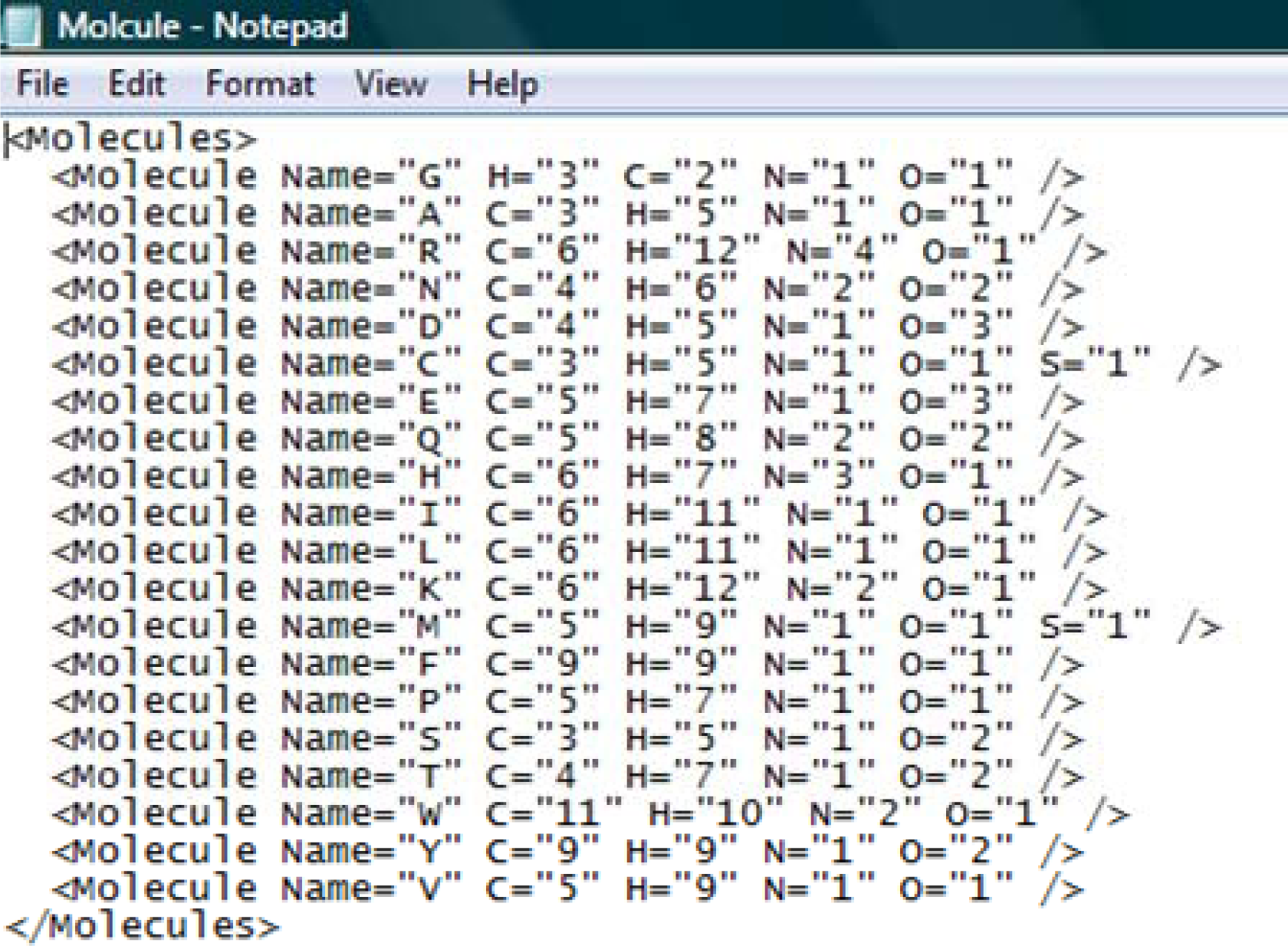
A portion of <u>Molecule.XML</u>.

**Figure 5.**
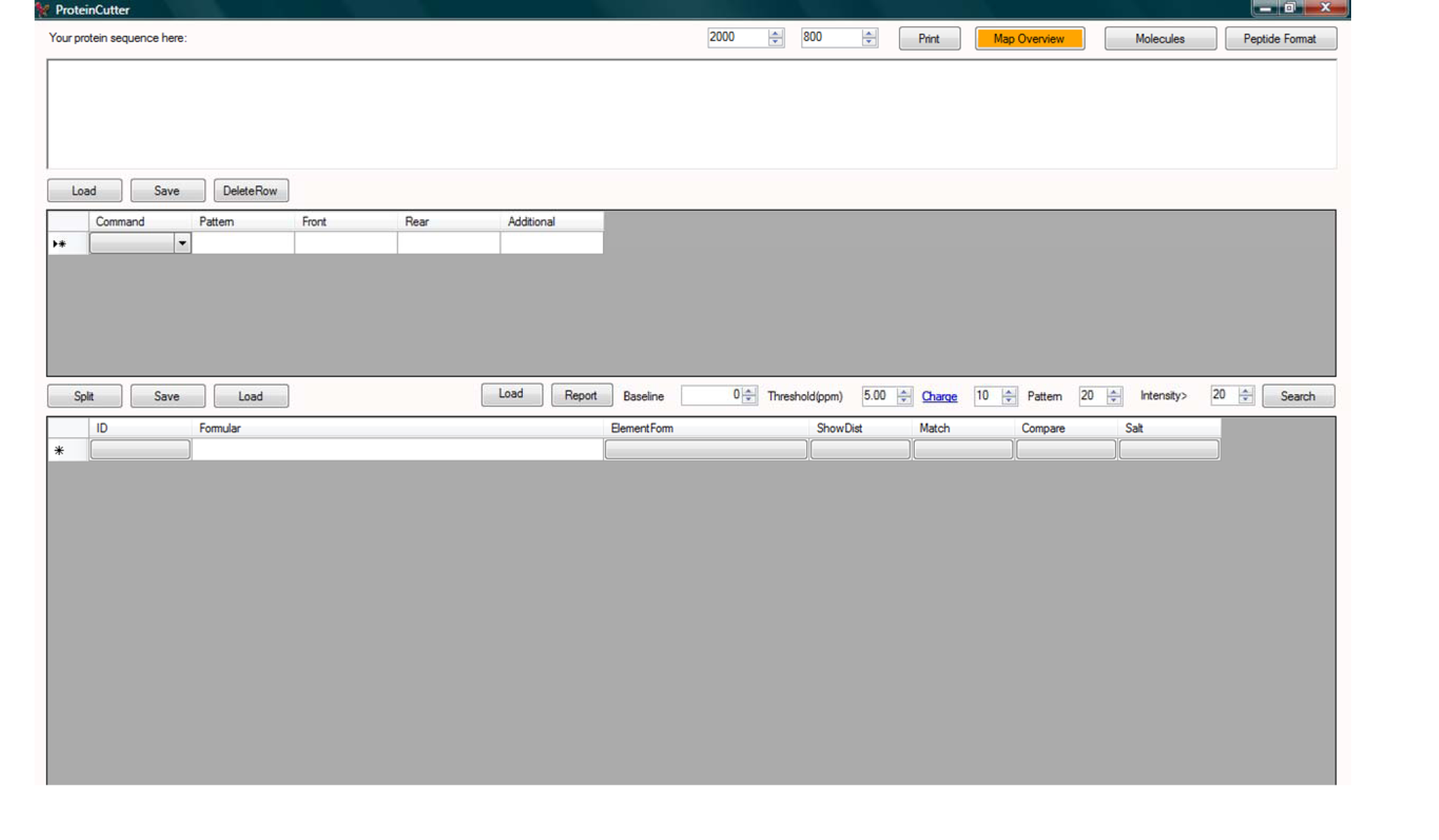
The full user-interface of PolyCut.

**Figure 6.**
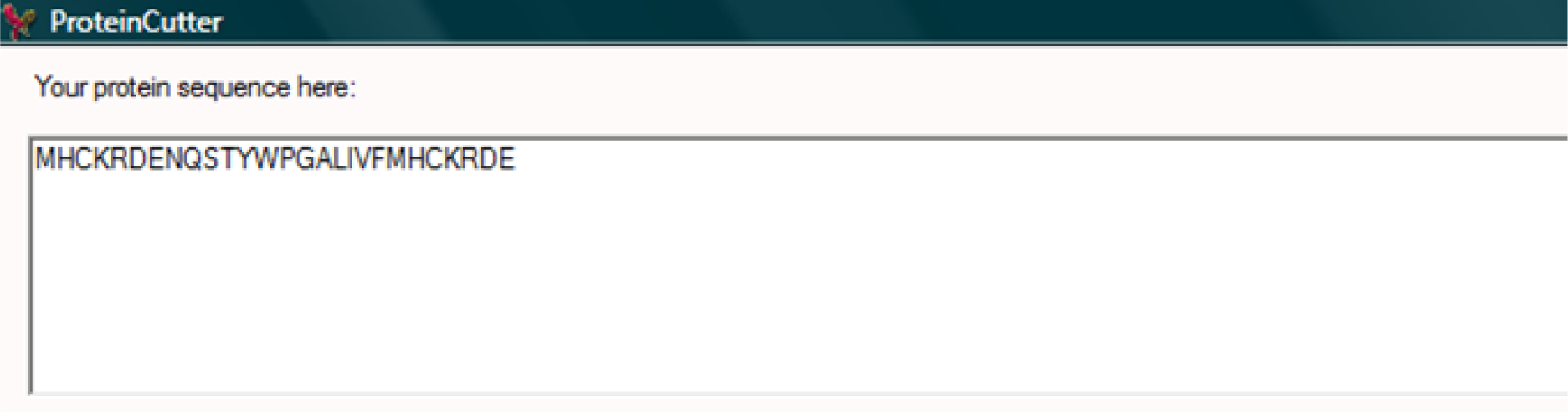
A portion of the user-interface of PolyCut showing the place for where a protein (or any polymer with residues already defined in figure 4) sequence is entered.

**Figure 7.**
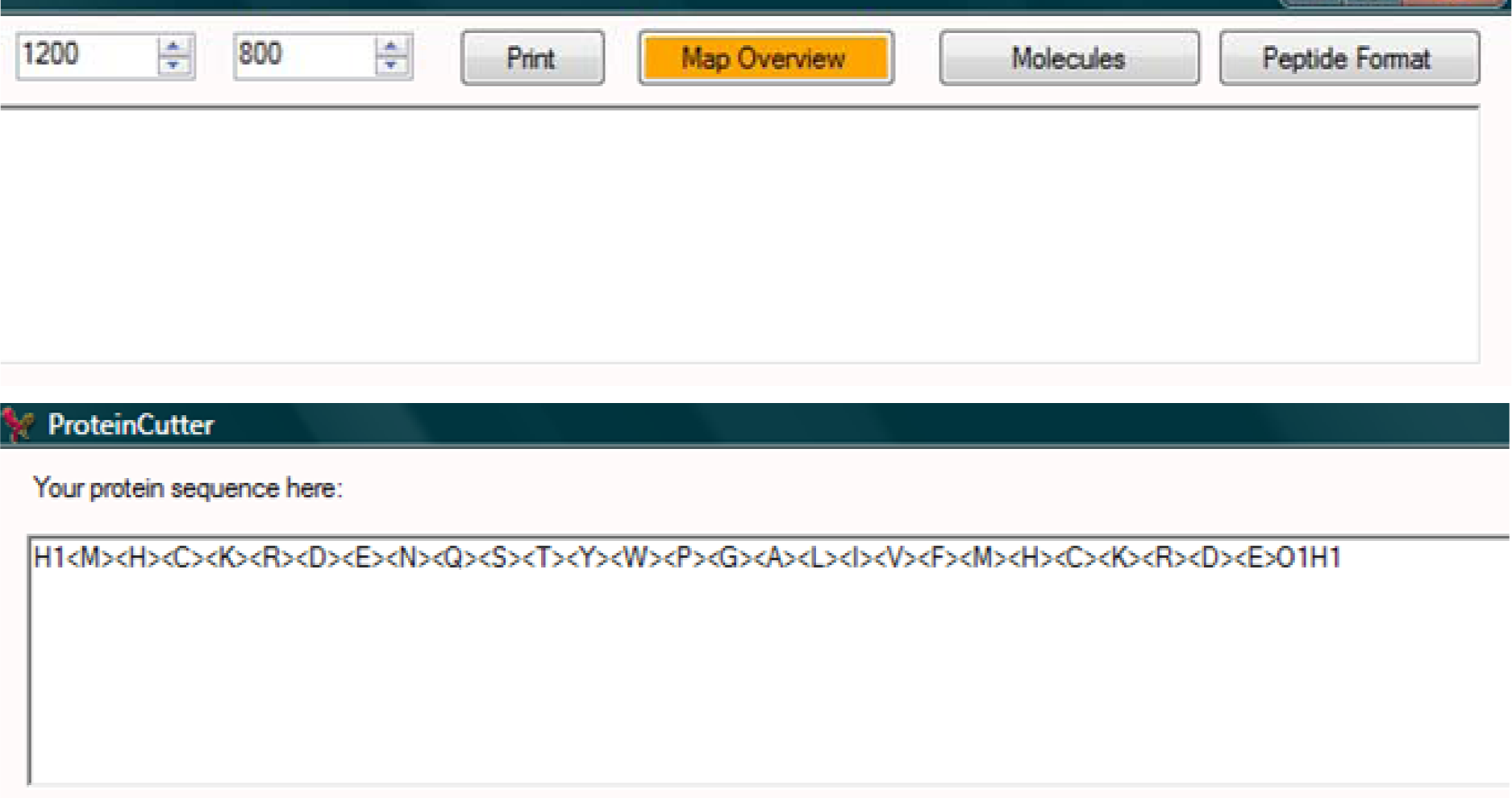
A portion of the user-interface of PolyCut. Clicking the button ‘Peptide Format’ changes the entered protein sequence (figure 6) to PolyCut-readable form. The N- and C-termini need to be defined. For example, a ‘H1′ at N-terminal and a ‘O1H1′ at C-terminal indicate that the sequence is an intact protein, a peptide from the hydrolysis at peptide bond(s) of a protein or a *y’* fragment from the fragmentation at a peptide bond of a protein via low-energy collisions. A ‘H1′ at N-terminal and a ‘H-1′ at C-terminal indicate that the sequence is a *b* fragment from the fragmentation at a peptide bond of a protein via low-energy collisions.

**Figure 8.**
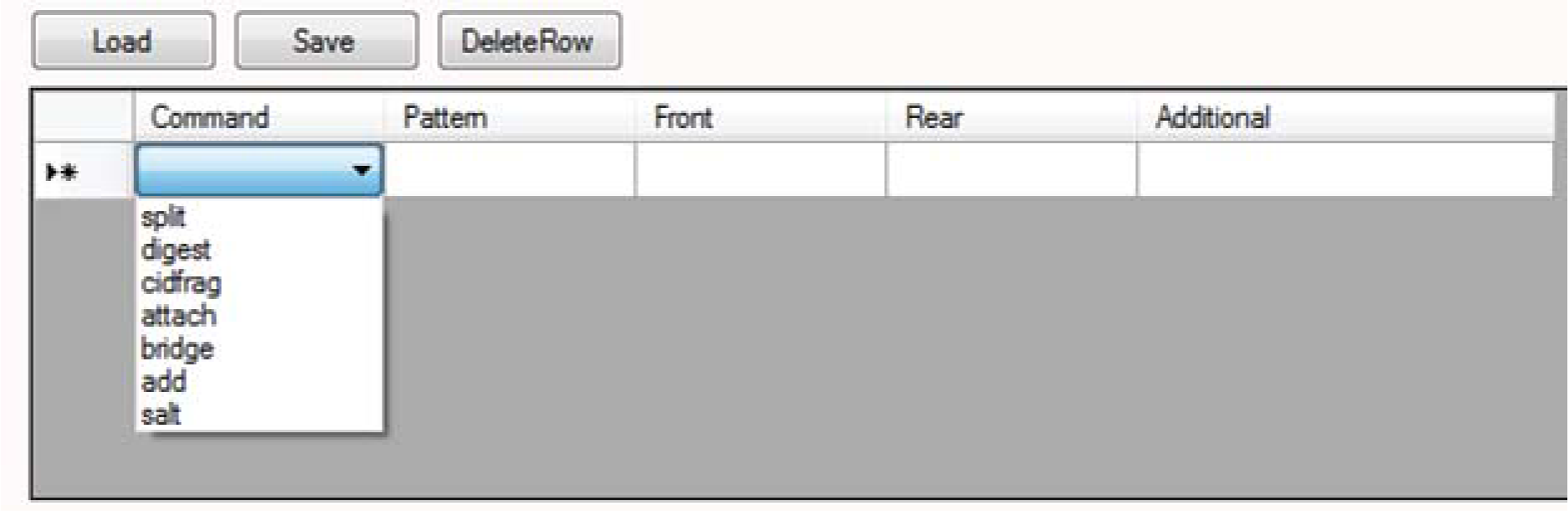
A portion of the user-interface of PolyCut to define the fragmentation or digestion site(s) on the entered sequence (figure 7) and the molecular structures of the resulting fragments or molecules.

**Figure 9.**
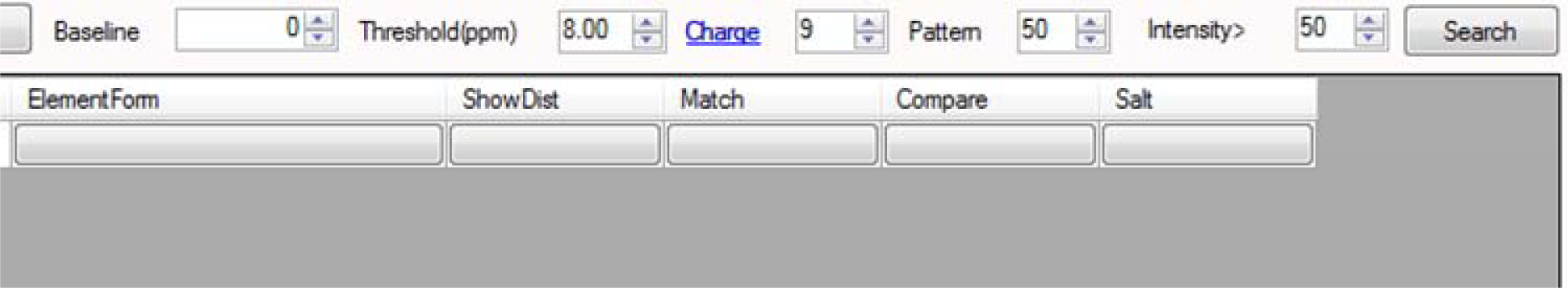
A portion of the user-interface of PolyCut which defines the highest possible charge state for the theoretical chemical species generated and the criteria for an ‘assignment match’.

**Figure 10.**
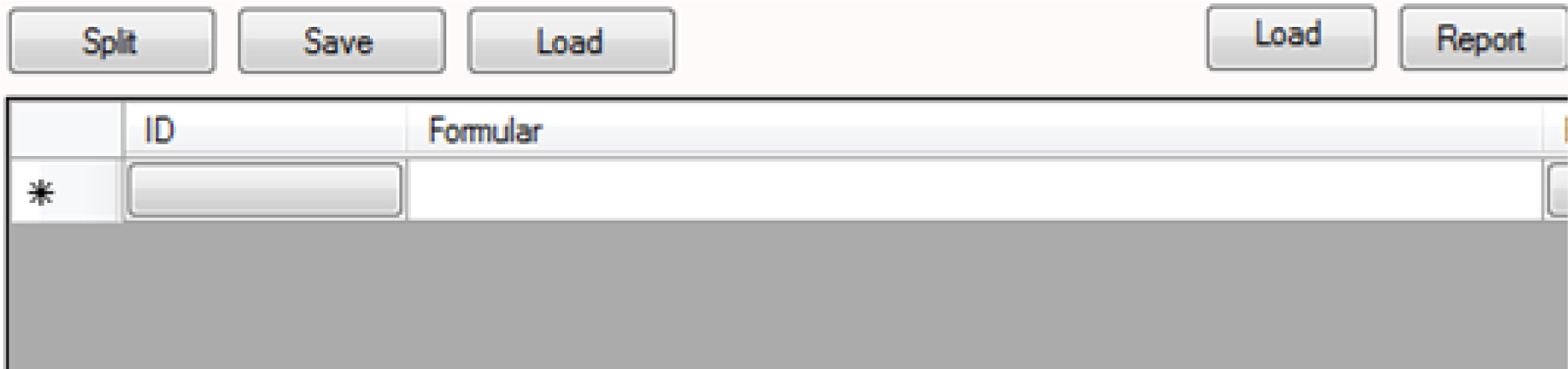
A portion of the user-interface of PolyCut which shows the list of theoretical chemical species.

**Figure 11.**
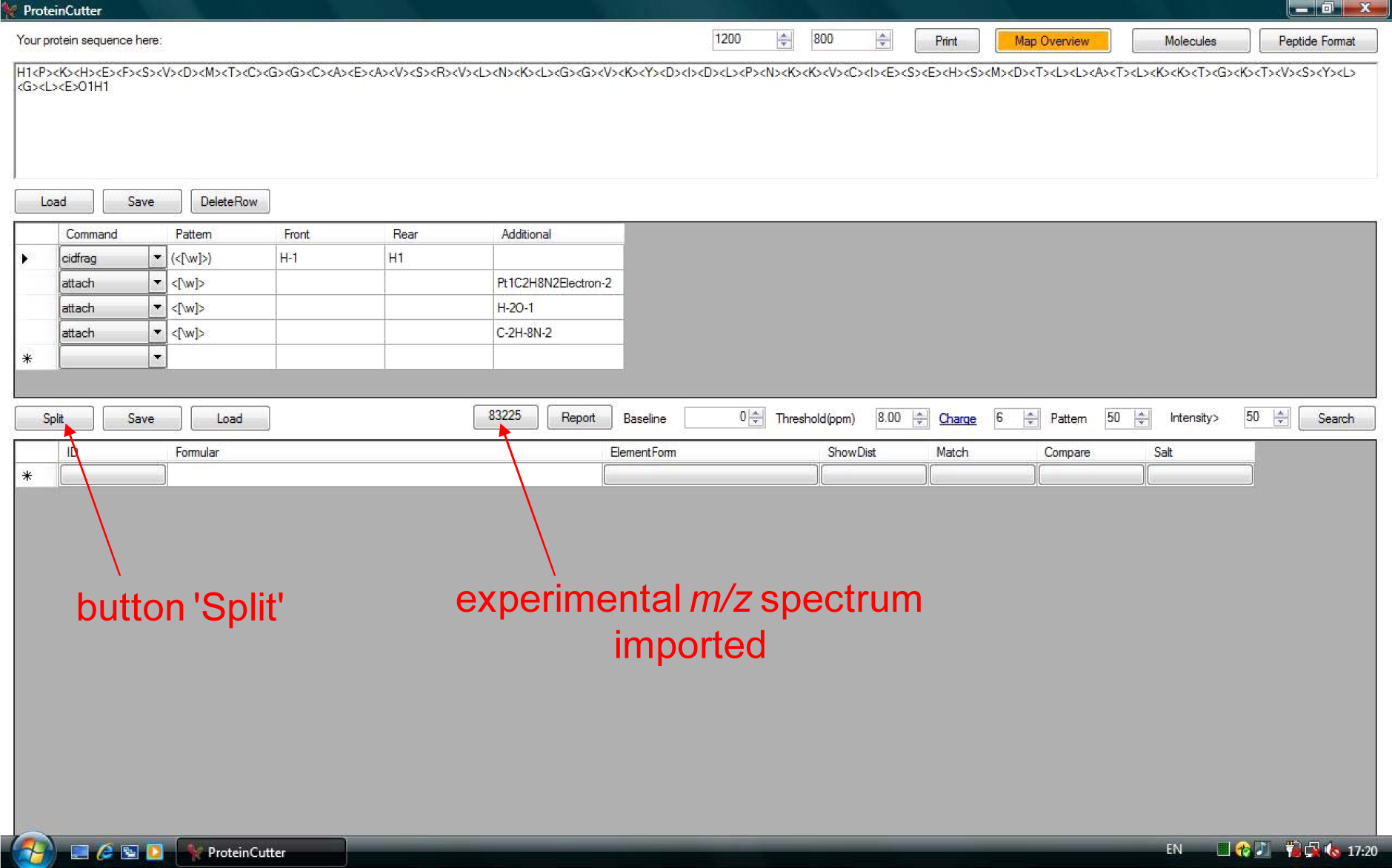
The user-interface of PolyCut.

**Figure 12.**
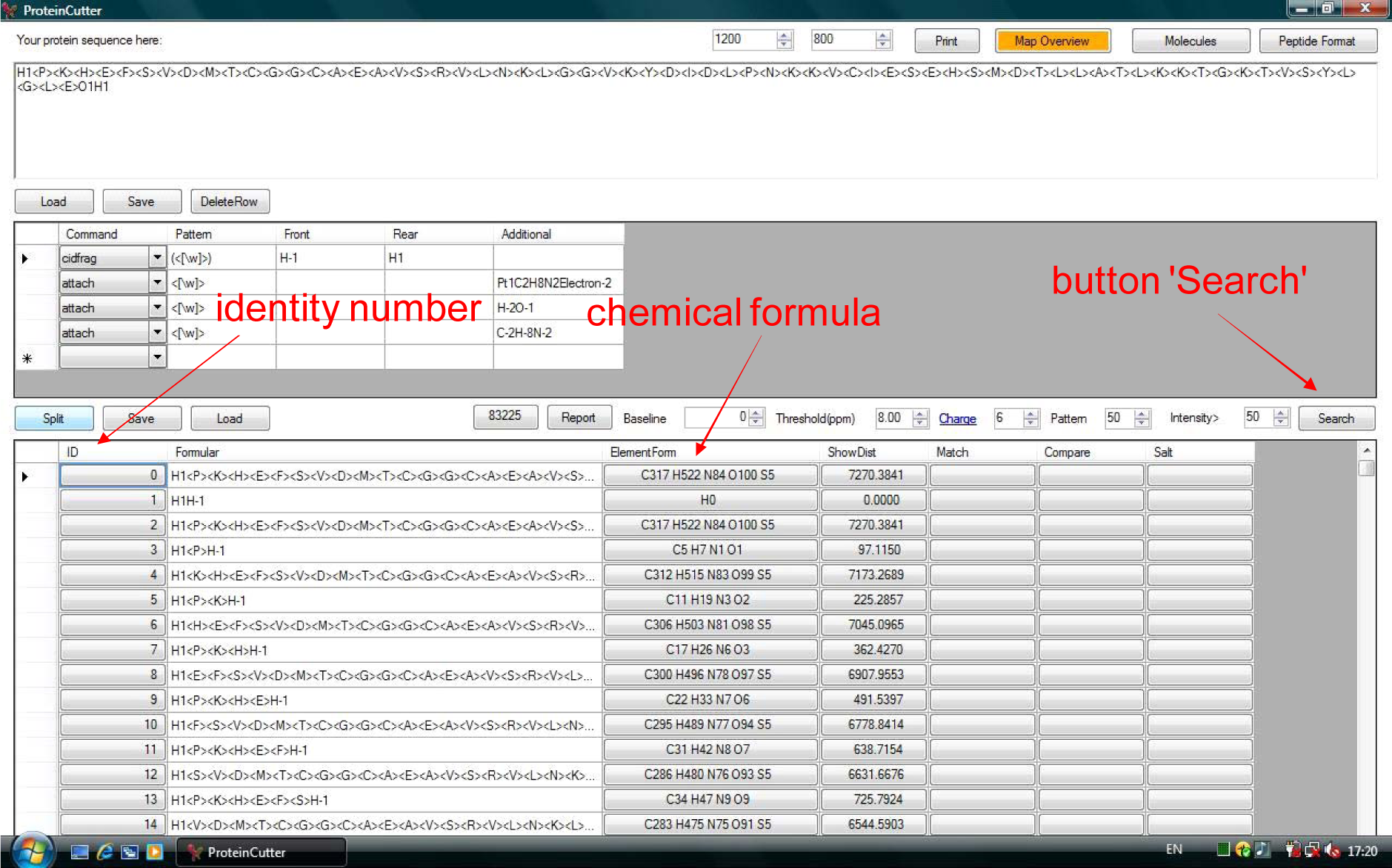
The user-interface of PolyCut.

**Figure 13.**
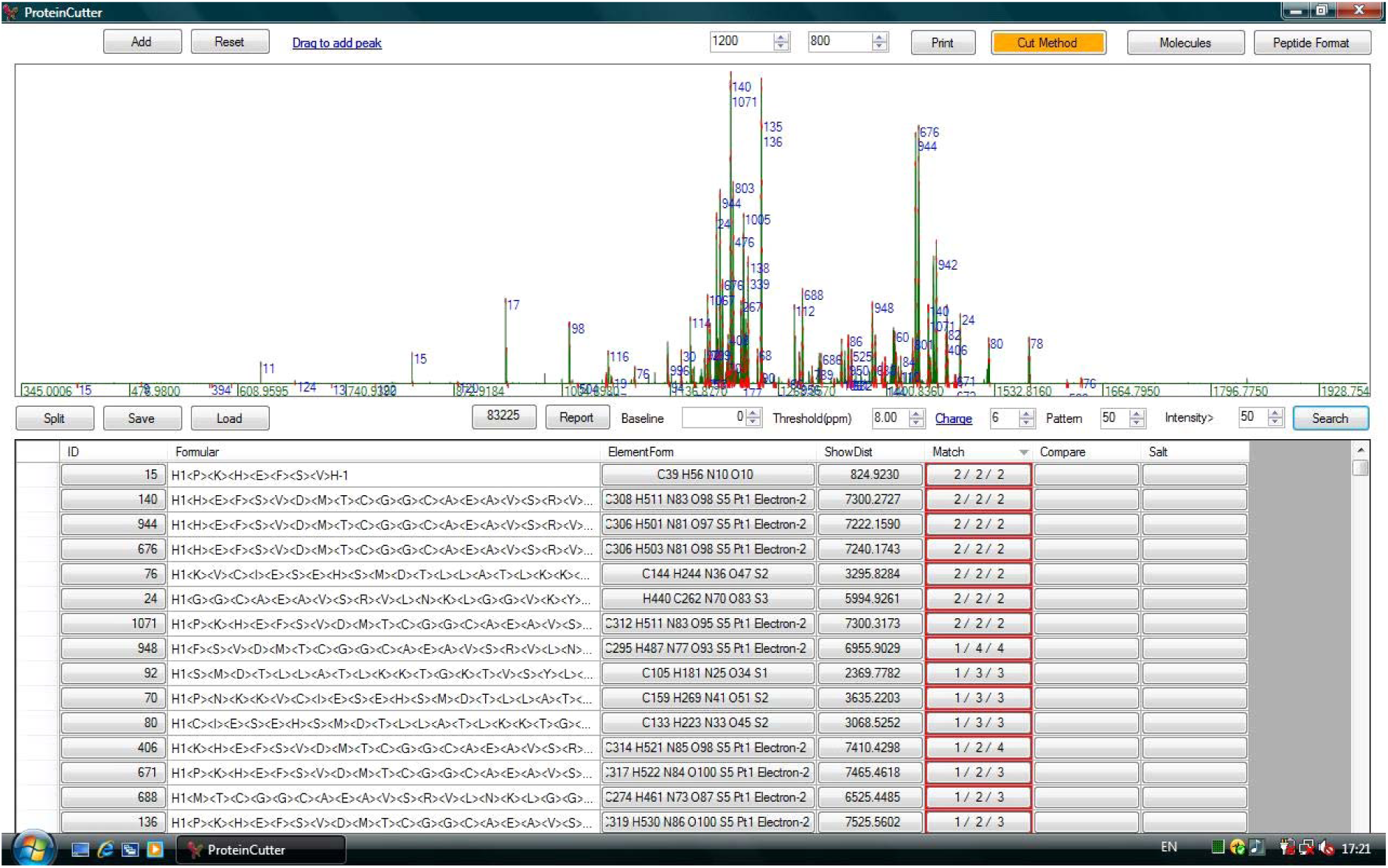
The user-interface of PolyCut.

**Figure 14.**
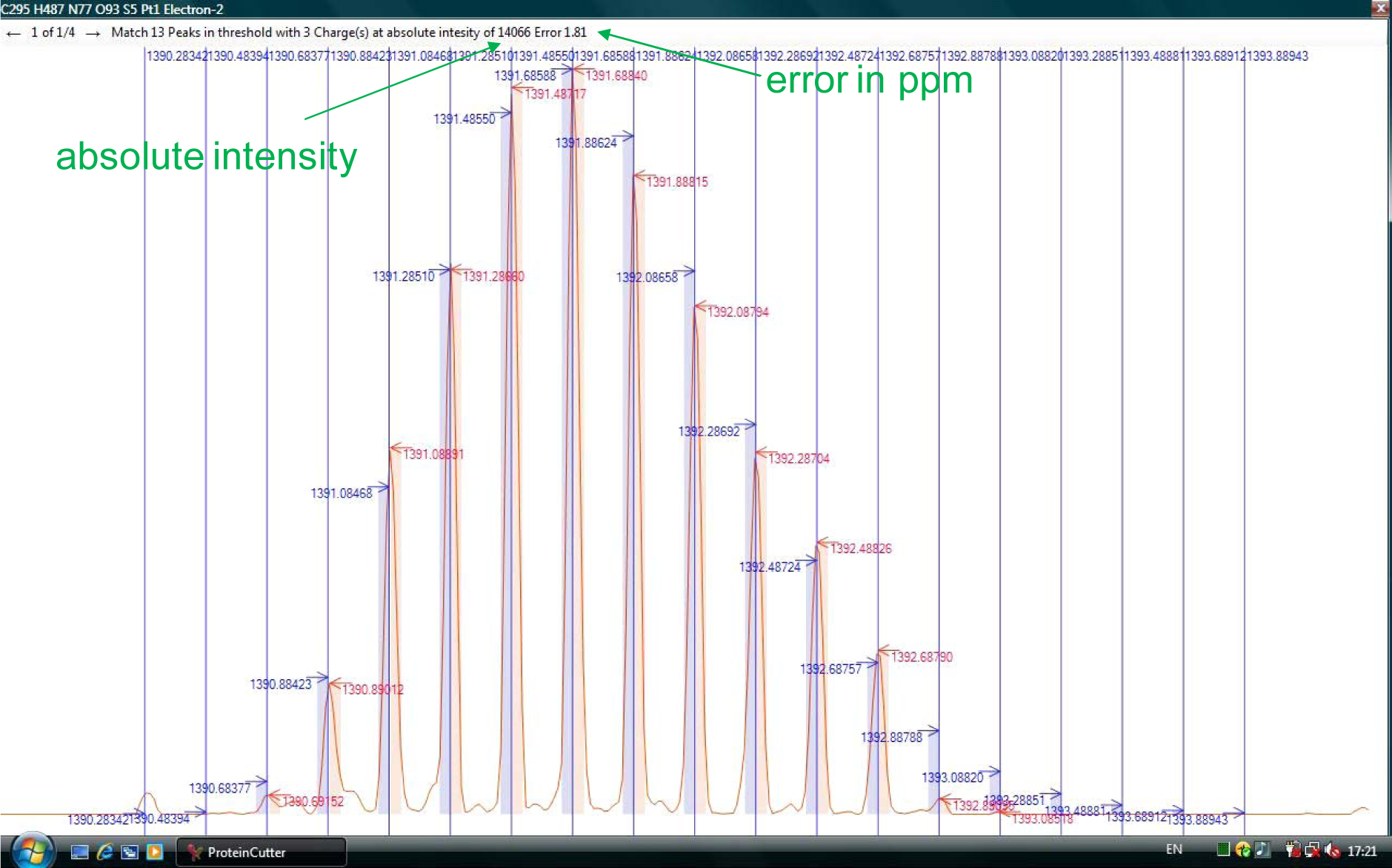
The user-interface of PolyCut.

## Author Contributions

[More].

Not to identify peptide nor protein, not for sequencing, but for identification of fragments or digested peptides from know sequence. Not for database.

## Appendix 1. Lists of MS softwares. [More].

### List A (softwares for assignments of PMF and/or MSn signals)

1. ProteinProspector v 5.6.2 (http://prospector2.ucsf.edu/prospector/mshome.htm).
2. ExPASy Proteomics Server (http://www.expasy.org/).
3. Search engines SEQUEST, Mascot, Sonar, MS-Tag (ProteinProspector) [*Anal*. *Chem*. **2002**, *74(20)*, 5383-5392; *Anal*. *Chem*. **2003**, *75(17)*, 4646-4658.].
4. X! Tandem.
5. PEAKS (*de novo* sequencing algorithm) [*Anal*. *Chem*. **2004**, *76(8)*, 2220-2230.].
6. SEQUEST, ProteinLynx (database-searching programmes) [*Anal*. *Chem*. **2004**, *76(8)*, 2220-2230.].
7. Guten Tag (database-searching programmes) [*Anal*. *Chem*. **2003**, *75(23)*, 6415-6421.].
8. OpenSea [*Anal*. *Chem*. **2004**, *76(8)*, 2220-2230.].
9. ProteinCalculator (Xcalibur 2.0 SR1, FT Programs 2.0.1.0611, Thermo Electron Corporation).
10. Bioworks and Proteome Discoverer 1.0 (Thermo Finnigan).
11. Spectrum Mill (Agilent).
12. Water Identity PLGS 2.4 (Waters).
13. Scaffold 3 (http://www.proteomesoftware.com/).
14. massXpert version 1.7.6 (http://www.massxpert.org). Read v 2.3.5 manual, try v 2.4.0.
15. Phenyx.
16. OMSSA.
17. THRASH (J. Am. Soc. Mass Spectrom. 2000, 11(4), 320-332.).
18. ICR-2LS.
19. MIDAS (http://magnet.fsu.edu/~midas).
20. STEM (*J*. *Proteome Res*. **2005**, *4(5)*, 1826-1831.). Check MS webpages in MSA, MSB…
21. http://www.proteomesoftware.com/, http://proteome-software.wikispaces.com/FAQ, http://proteome-software.wikispaces.com/FAQ+Complete#Complete List of FAQs, http://au.expasy.org/sprot/, http://au.expasy.org/tools/).
22. ProteinCutter (http://biochemie.upol.cz/software/proteincutter/, http://citimseskvele.eu/bch/).
23. More? (See Scaffold 3 web).
24. MS data format (http://en.wikipedia.org/wiki/Mass_spectrometry_data_formats).
25. MS software (http://en.wikipedia.org/wiki/Mass_spectrometry_software).
26. More?
27. LutefiskXP (http://www.hairyfatguy.com/lutefisk/)
28. Sherpa (http://www.hairyfatguy.com/Sherpa/)

Also the software list [*Biochim*. *Biophys*. *Acta*-*Proteins Proteomics* **2003**, *1646(1-2)*, 1-10.]. De-isotope: [A. R. Ledvina, M. M. Savitski, A. R. Zubarev, D. M. Good, J. J. Coon and R. A. Zubarev, *Anal*. *Chem*. **2011**, DOI: 10.1021/ac201843e.].

[*Anal*. *Chem*. **1994**, *66(24)*, 4390-4399; *Proc*. *Natl*. *Acad*. *Sci*. *U*. *S*. *A*. **1996**, *93(16)*, 8264-8267; *J*. *Am*. *Soc*. *Mass Spectrom*. **2005**, *16(12)*, 2027-2038.].

Require recognisable mono-isotopes by looking at mono-isotope first to generate molecular formula, then theoretical isotope pattern against experimental isotope pattern, at least 3 isotope peaks are present then consider a match, examples of < 1000 Da, Ru is the only heavy element, not for MSn, different purpose: to guess formulae then eliminate the worst ones, does not consider Ru as a fixed charge [*J*. *Am*. *Soc*. *Mass Spectrom*. **2006**, *17(12)*, 1692-1699.].

During our software development, we noticed a recent software publication for theoretical isotope pattern against experimental isotope pattern for HDX experiments:

[Z-Y Kan, L. Mayne, P. S. Chetty and S. W. Englander, *J*. *Am*. *Soc*. *Mass Spectrom*. **2011**, DOI: 10.1007/s13361-011-0236-3.].

Cross-link analyses including the disulfide bond [M. Götze, J. Pettelkau, S. Schaks, K. Bosse, C. H. Ihling, F. Krauth, R. Fritzsche, U. Kühn and A. Sinz, *J*. *Am*. Soc. Mass *Spectrom*. **2011**, DOI: 10.1007/s13361-011-0261-2.].

[D. E. Miller, C. B. Prasannan, M. T. Villar, A. W. Fenton and A. Artigues, *J*. *Am*. *Soc*. *Mass Spectrom*. **2011**, DOI: 10.1007/s13361-011-0234-5.].

Software for SILAC [*Anal*. *Chem*. **2011**, *83(22)*, 8403-8410.].

This programme is designed for mass spectral data acquired with data independent acquisition. Mass spectral data from data dependent acquisition do not work on this programme.

Do not do deconvolution to remain charge state info. Tell you which charge state of which fragment is found. Charge remaining on fragment provides clues on fragmentation mechnaisms.

Can be applied to oligopeptide, oligonucleotide, oligosaccharide, lipid, PNA, glycopeptide, glycolipid, peptoid.

Emphasize in introduction all PolyCut function: why made them?

PolyCut not for sequencing. MSn spectra contains no sequencial information anyway.

To best of knowledge, this is the first programme handling low mono- transition metal with oxidation states, for metallomics and metalloproteomics.

Not necessarily following the natural abundance. Designed for working offline.

Mechanism scanner.

Distinguish two or more overlapping isotope patterns of individual species.

Can behave as simulator for isotope patterns also for various MS1 ions, MSn ions, with recognition of fixed charges.

[S. M. Miladinovic, A. N. Kozhinov, M. V. Gorshkov and Y. O. Tsybin, *Anal*. *Chem*., **2012**, DOI: 10.1021/ac2034584.].

### List B (softwares for simulations of isotope patterns)

1. Qual Browser version 2.0 (Thermo Electron Corporation).
2. Isotope Distribution Calculator version 3.1.346.14 (Agilent).

### List C (resources for post-translational modifications)

1. ABRF Delta Mass (http://www.abrf.org/index.cfm/dm.home?AvgMass=all).
2. RESID Database (http://home.earthlink.net/~jsgaravelli/RESIDInfo.HTML).
3. UNIMOD.

## Appendix 2.

Figures A2.1 to A2.16 are examples of generating different lists of theoretical chemical species. Theoretical isotope patterns for different charge states are simulated and overlaid against the imported experimental *m/z* spectrum.

**Figure A2.1.**
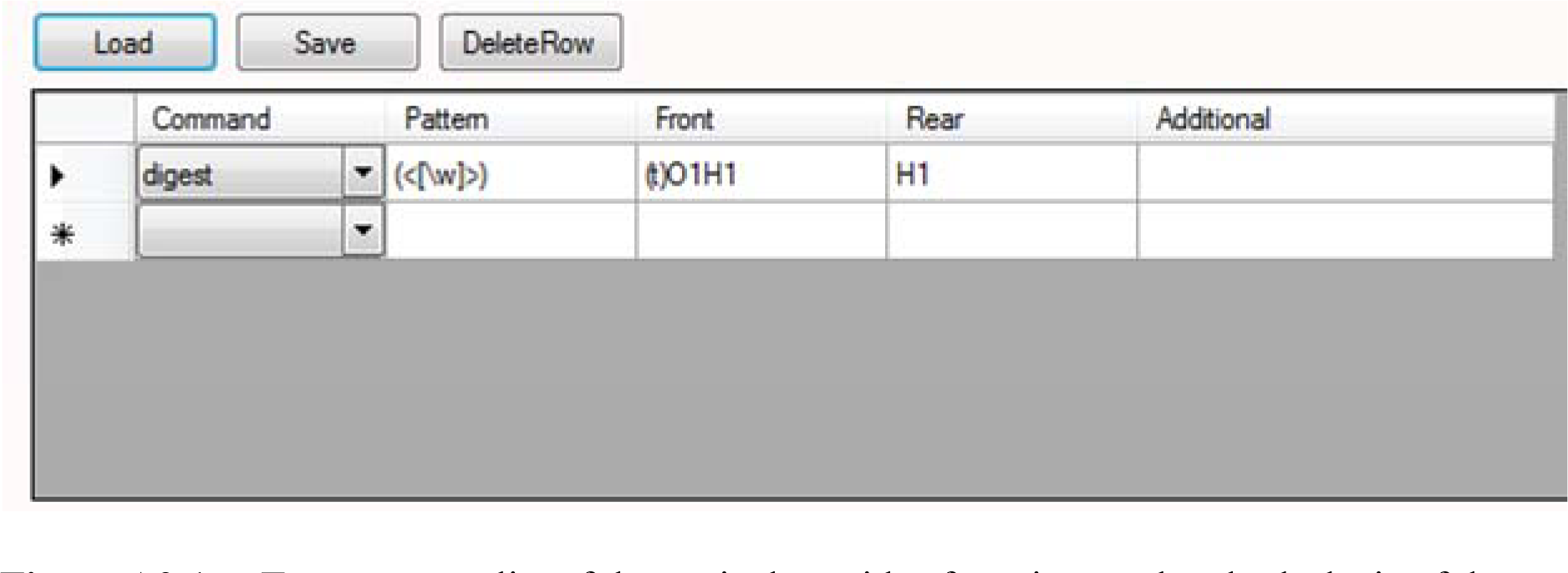
To generate a list of theoretical peptides from incomplete hydrolysis of the entered protein sequence (figure 3.7) at randem peptide bonds.

**Figure A2.2.**
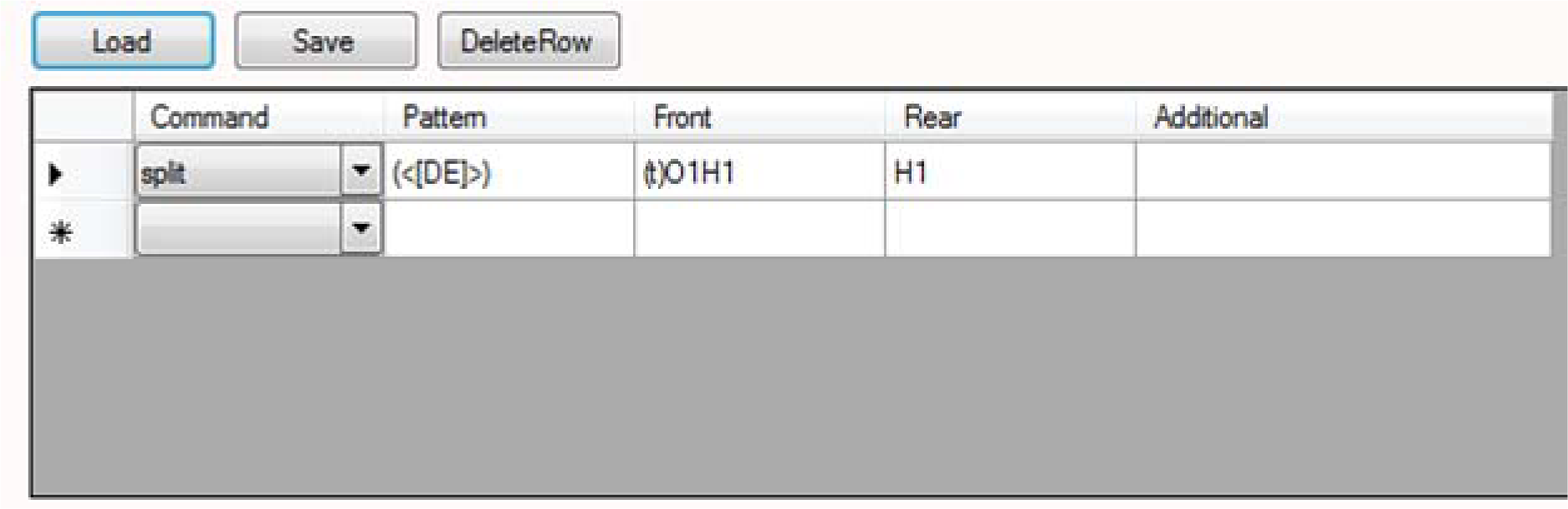
To generate a list of theoretical peptides from complete hydrolysis of the entered protein sequence (figure 3.7) at peptide bonds on the C-terminal side of Asp <u>and</u> Glu residues.

**Figure A2.3.**
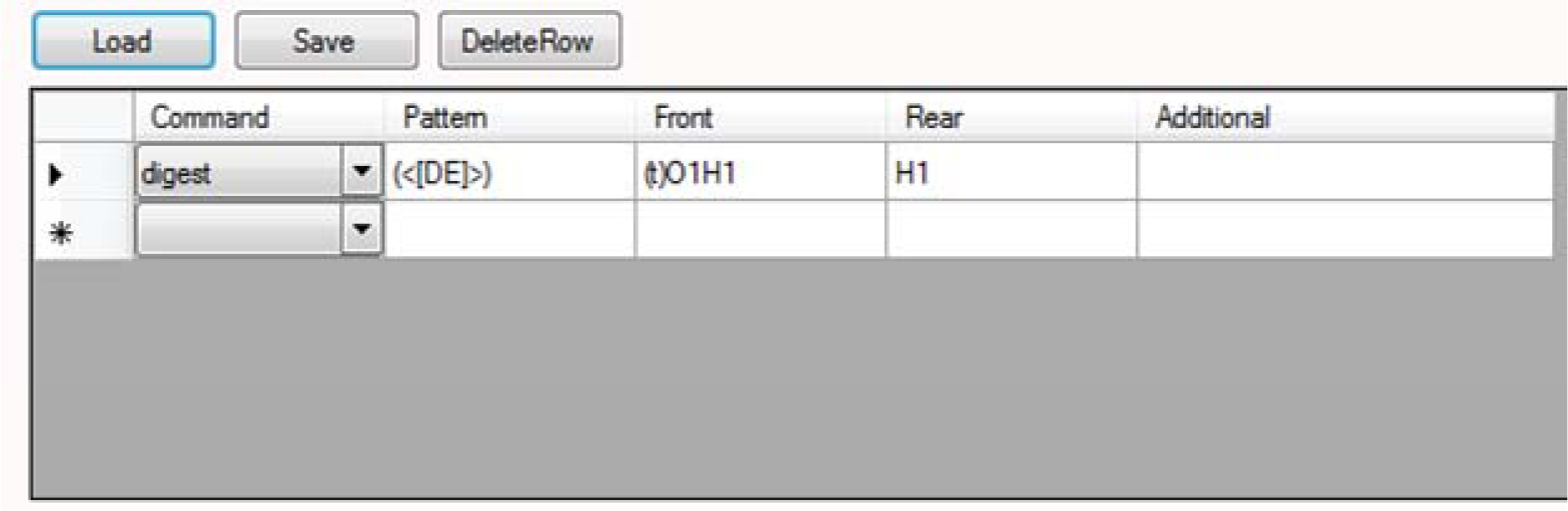
To generate a list of theoretical peptides from incomplete hydrolysis of the entered protein sequence (figure 3.7) at the peptide bonds on the C-terminal side of Asp <u>or</u> Glu residues.

**Figure A2.4.**
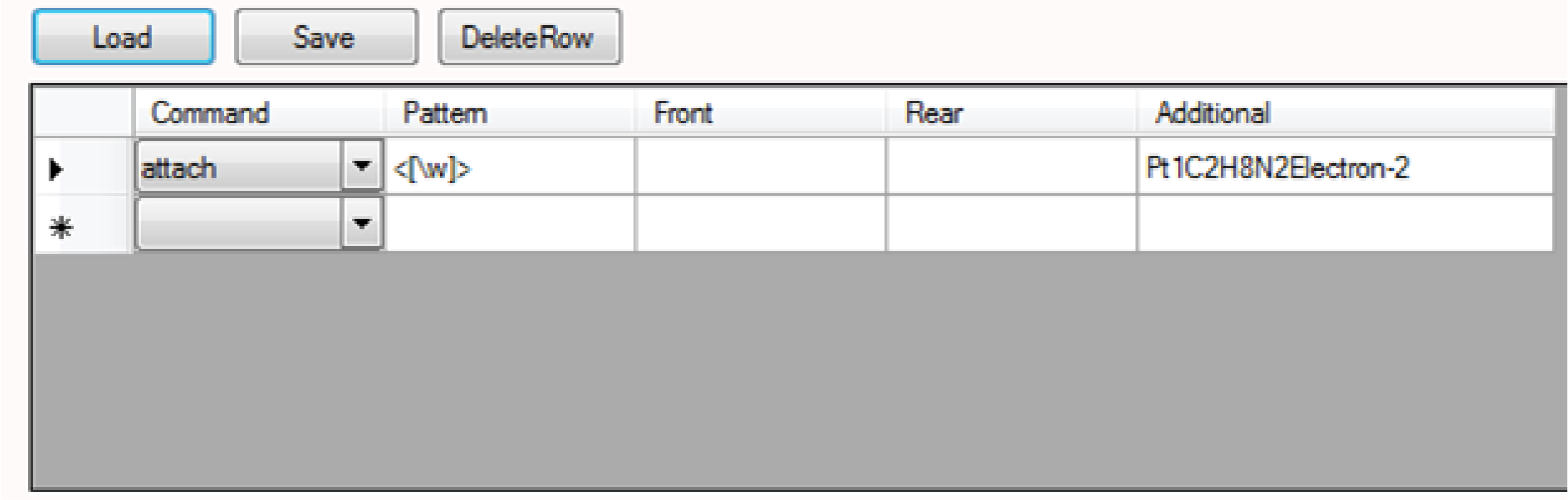
To generate a list of an theoretical intact protein with or without a Pt(en)^2+^ centre at any of the residues. ‘Electron-2’ is recognised by PolyCut to have a loss two-electron mass and have a +2 fixed charge prior to protonation or deprotonation while ‘Electron2′ is recognised to have a gain of two-electron mass and have a +2 fixed charge prior to protonation or deprotonation.

**Figure A2.5.**
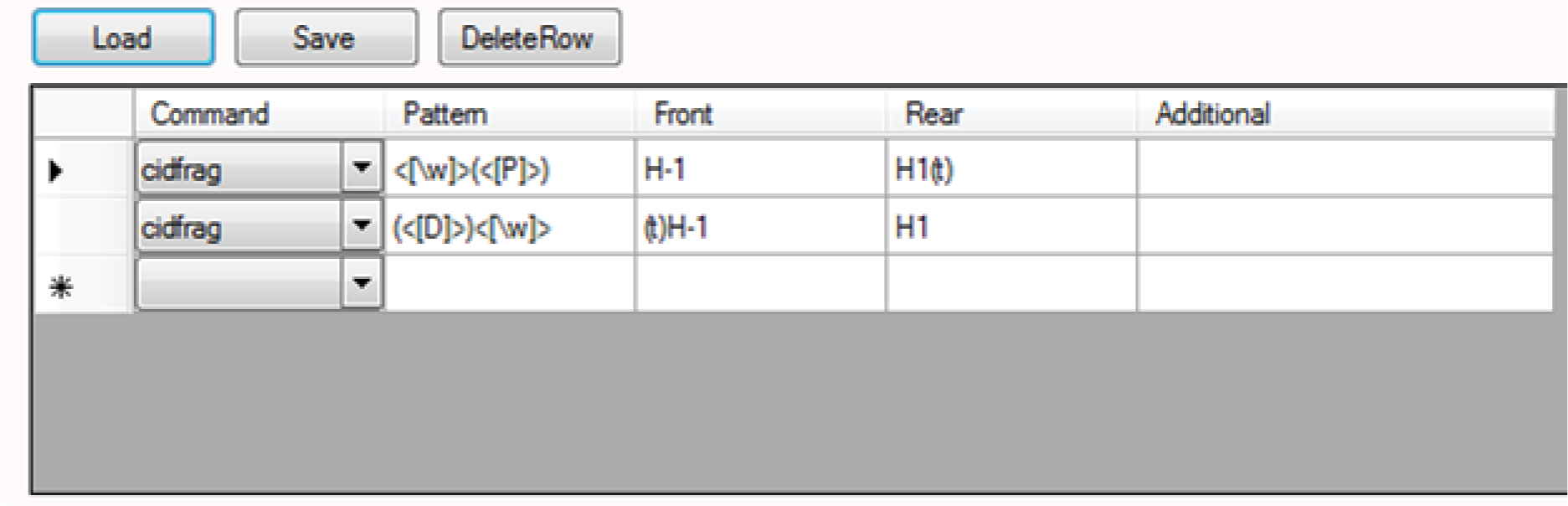
To generate a list of theoretical *b*/*y’* sequence pairs from a low-energy CID experiment of the entered protein sequence (figure 3.7). The fragment pairs arise from a single fragmentation at a peptide bond of the whole sequence on the N-terminal side of a Pro residue <u>or</u> C-terminal side of an Asp residue.

**Figure A2.6.**
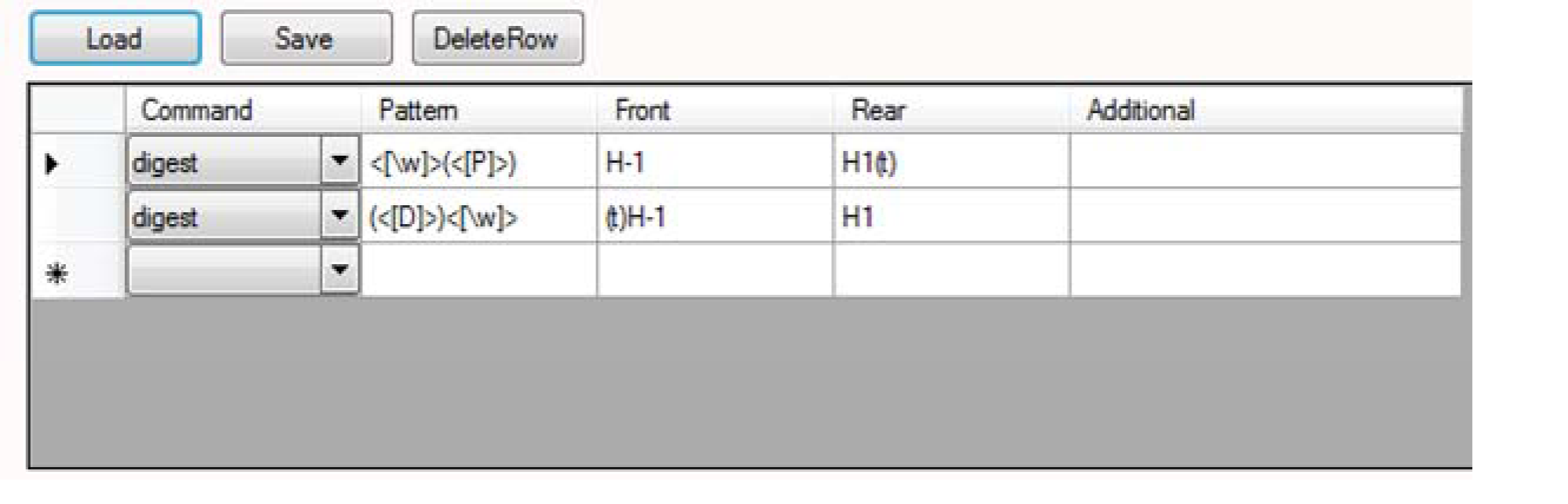
To generate a list of theoretical internal fragments for a low-energy CID experiment of the entered protein sequence (figure 3.7). The fragments arise from fragmentations of multiple peptide bonds of the whole sequence on the N-terminal side of Pro residues <u>and</u> C-terminal side of Asp residues.

**Figure A2.7.**
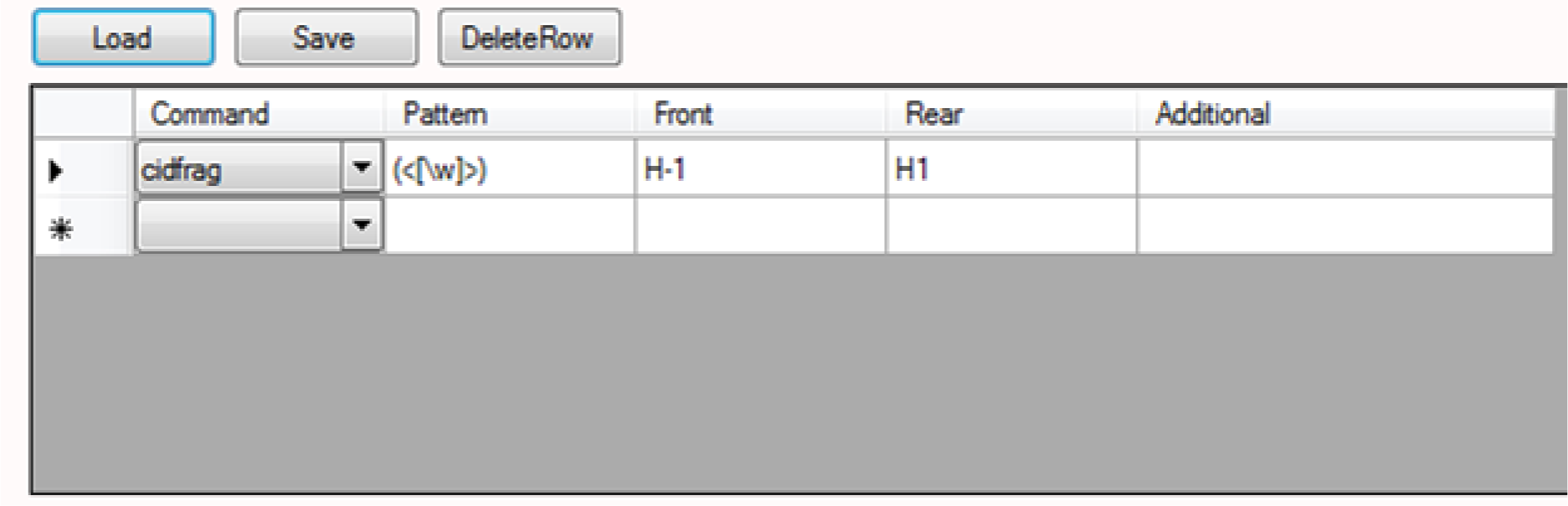
To generate a list of theoretical *b*/*y’* sequence pairs for a low-energy CID experiment of the entered protein sequence (figure 3.7). The fragment pairs arise from a single fragmentation at a randem peptide bond of the whole sequence.

**Figure A2.8.**
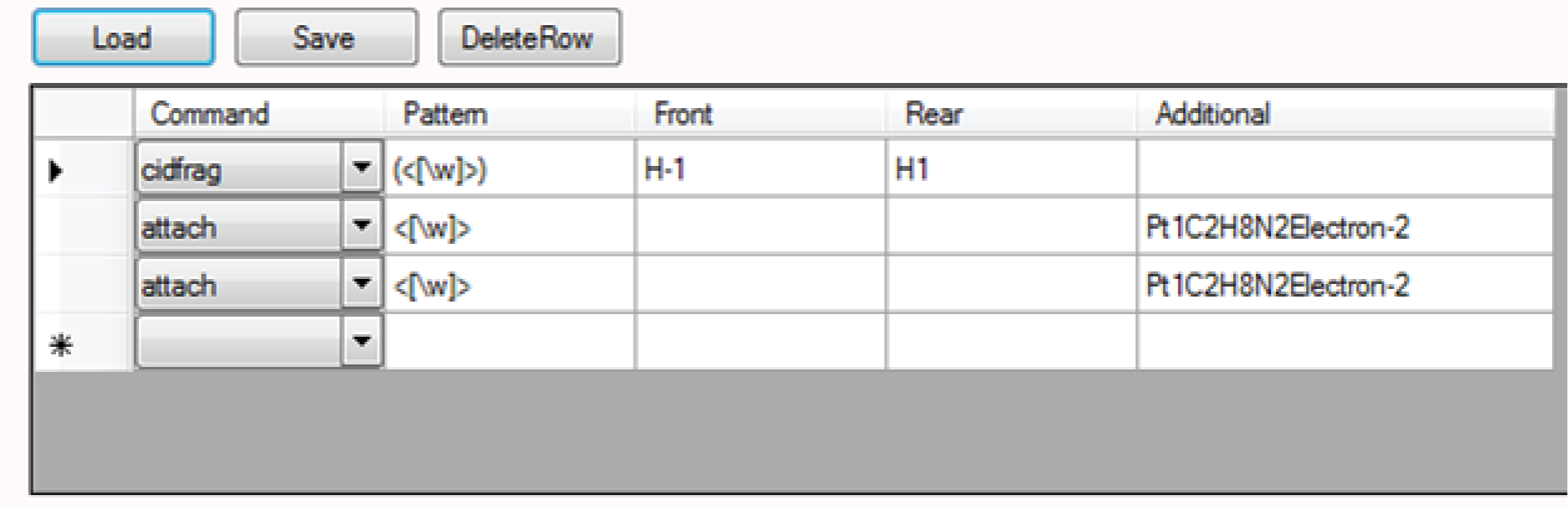
To generate a list of theoretical *b*/*y’* sequence pairs for a low-energy CID experiment of the entered protein sequence (figure 3.7). The fragment pairs arise from a single fragmentation at a randem peptide bond of the whole sequence with zero, one or two Pt(en)^2+^ centres on one of the residues.

**Figure A2.9.**
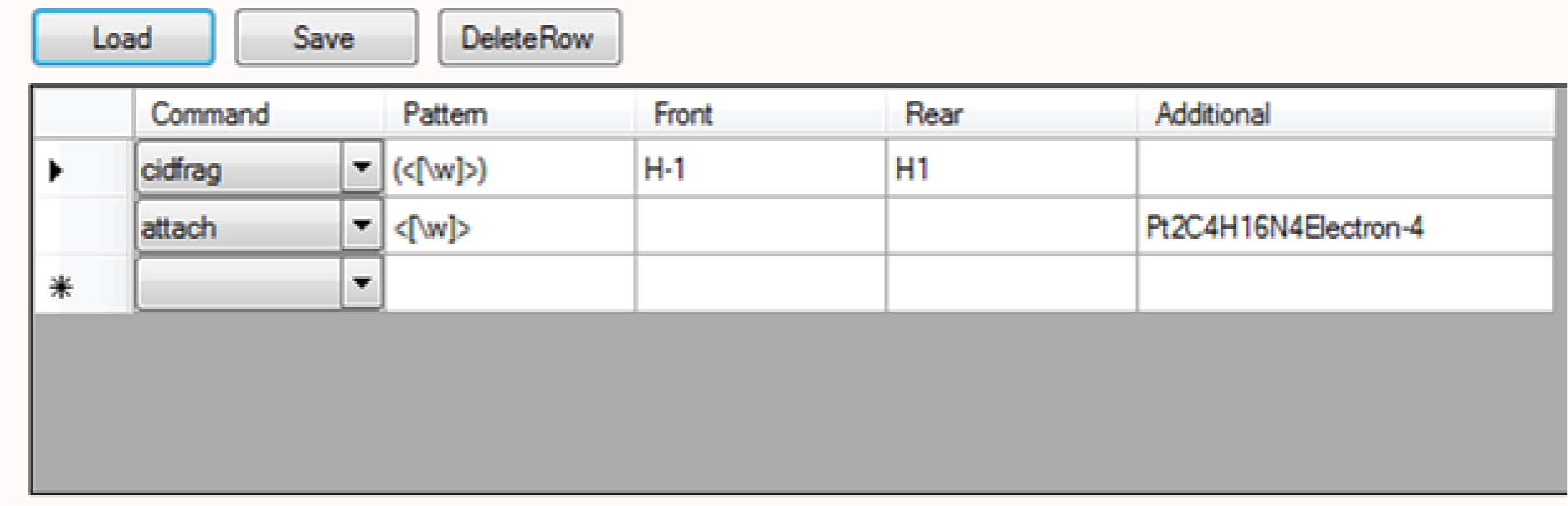
To generate a list of theoretical *b*/*y’* sequence pairs for a low-energy CID experiment of the entered protein sequence (figure 3.7). The fragment pairs arise from a single fragmentation at a randem peptide bond of the whole sequence with zero or two Pt(en)^2+^ centres on one of the residues.

**Figure A2.10.**
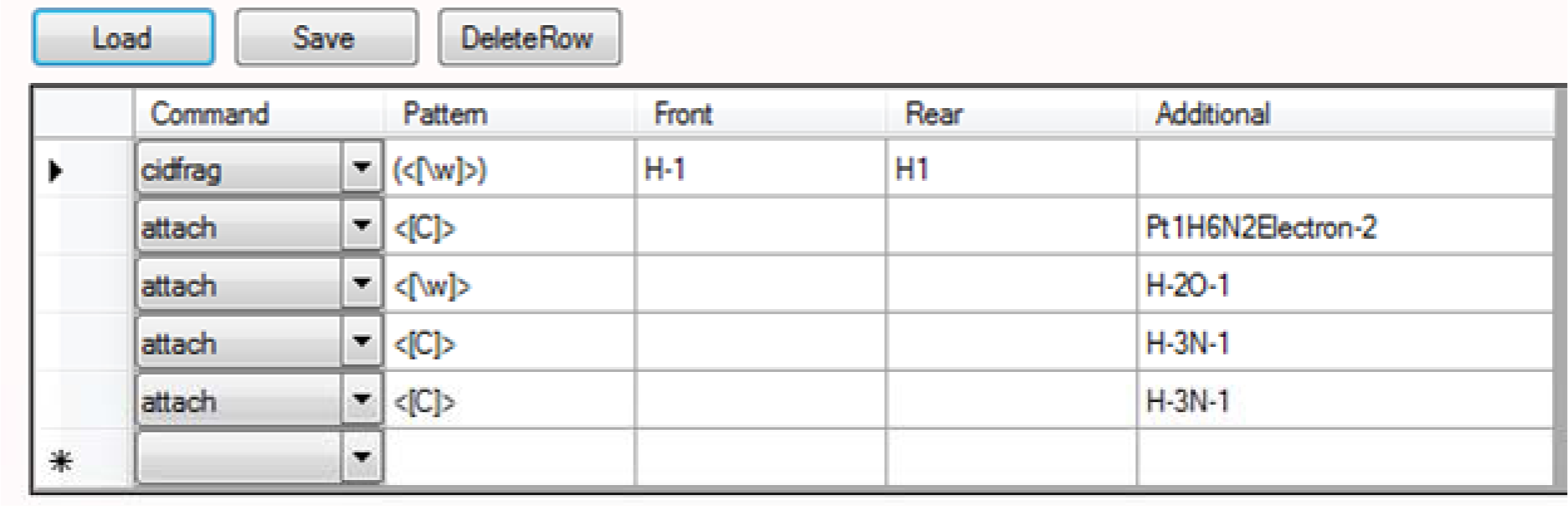
To generate a list of theoretical *b*/*y’* sequence pairs for a low-energy CID experiment of the entered protein sequence (figure 3.7). The fragment pairs arise from a single fragmentation of a randem peptide bond of the whole sequence, (i) with zero or one Pt(NH_3_)_2_^2+^ centre at one of the Cys residues, (ii) with zero or one neutral loss of water from one of the residues, and (iii) with zero, one or two neutral losses of ammonia from one or two of the Cys residues.

**Figure A2.11.**
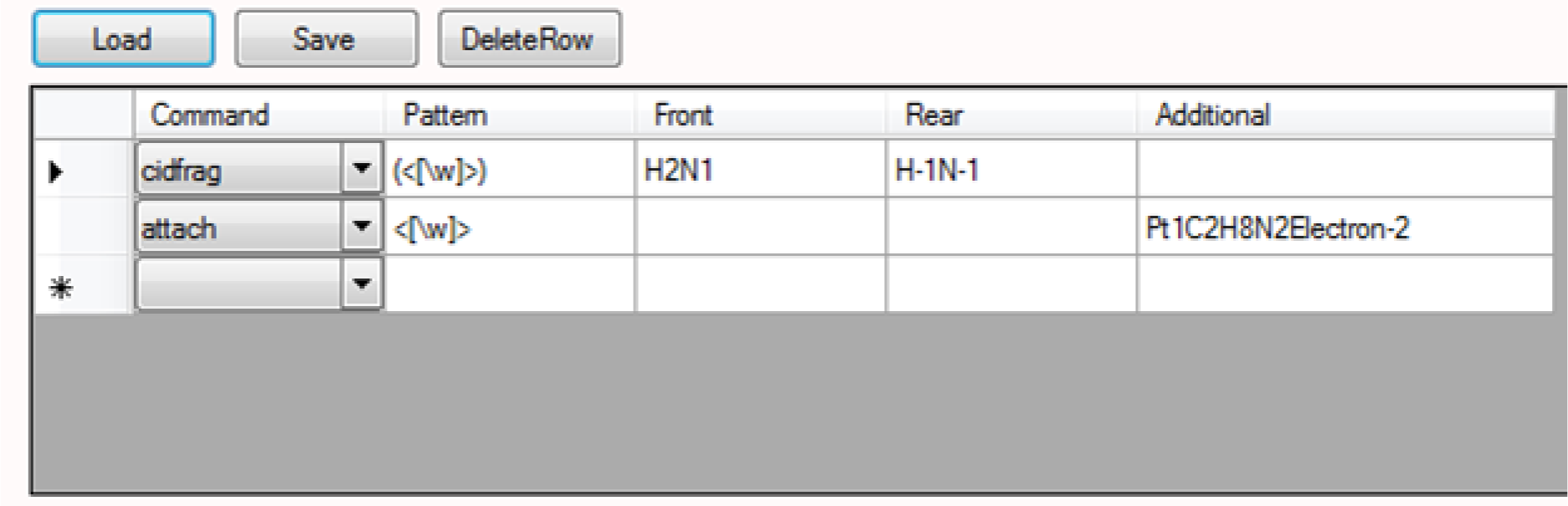
To generate a list of theoretical *c’*/*z*• sequence pairs for a ECD experiment of the entered protein sequence (figure 3.7). The fragment pairs arise from a single fragmentation of a random ψ bond on the whole sequence with zero or one Pt(en)^2+^ centre on one of the residues. Use your knowledge, PC is able to define *a*, *a*•, *a’*, and *b*-, *c*-, *x*-, *y*- and *z*-type (any other) ions.

**Figure A2.12.**
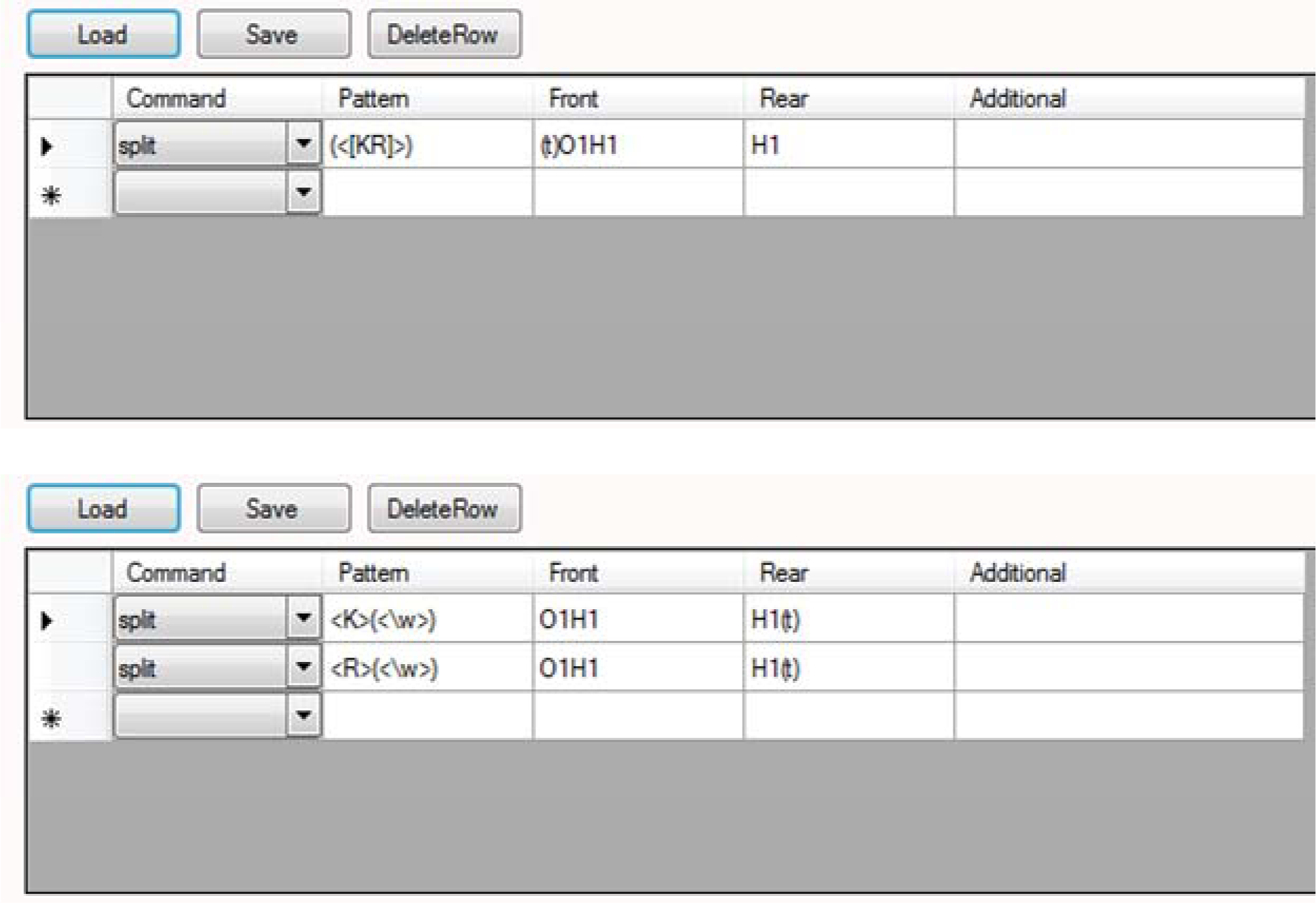
To generate a list of theoretical peptides from complete hydrolysis of the entered protein sequence (figure 3.7) at peptide bonds on the C-terminal side of Lys <u>and</u> Arg residues. The commands in the two panels are different but both lead to an identical list.

**Figure A2.13.**
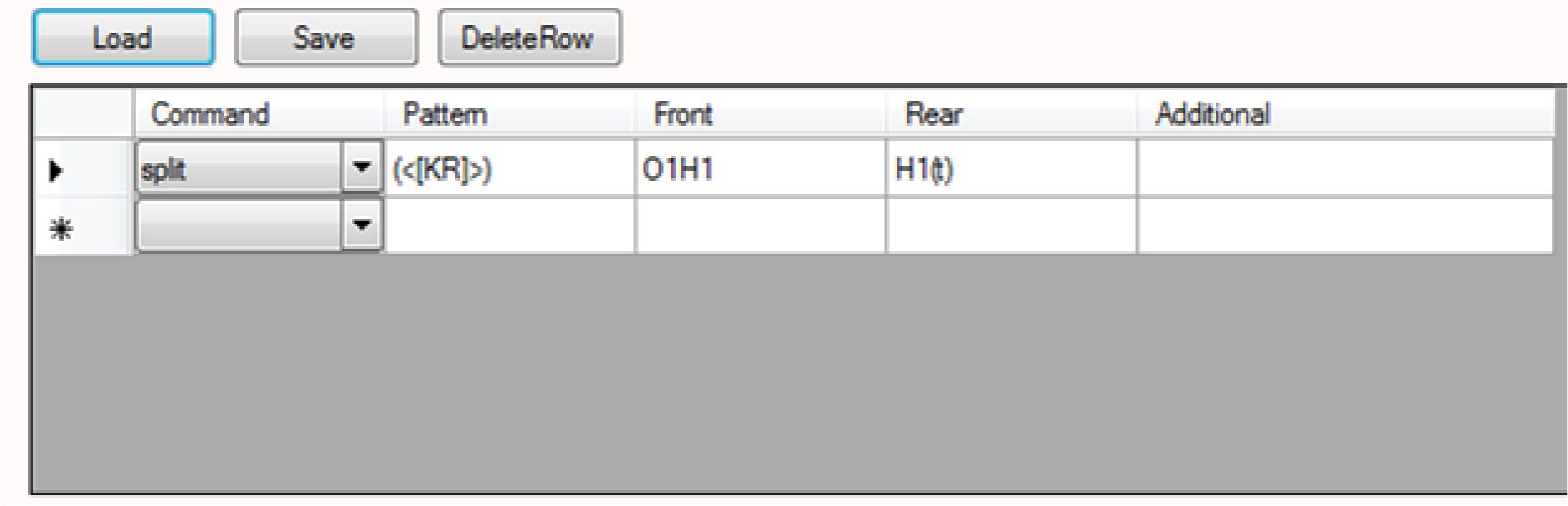
To generate a list of theoretical peptides from complete hydrolysis of the entered protein sequence (figure 3.7) at peptide bonds on the N-terminal side of Lys <u>and</u> Arg residues.

**Figure A2.14.**
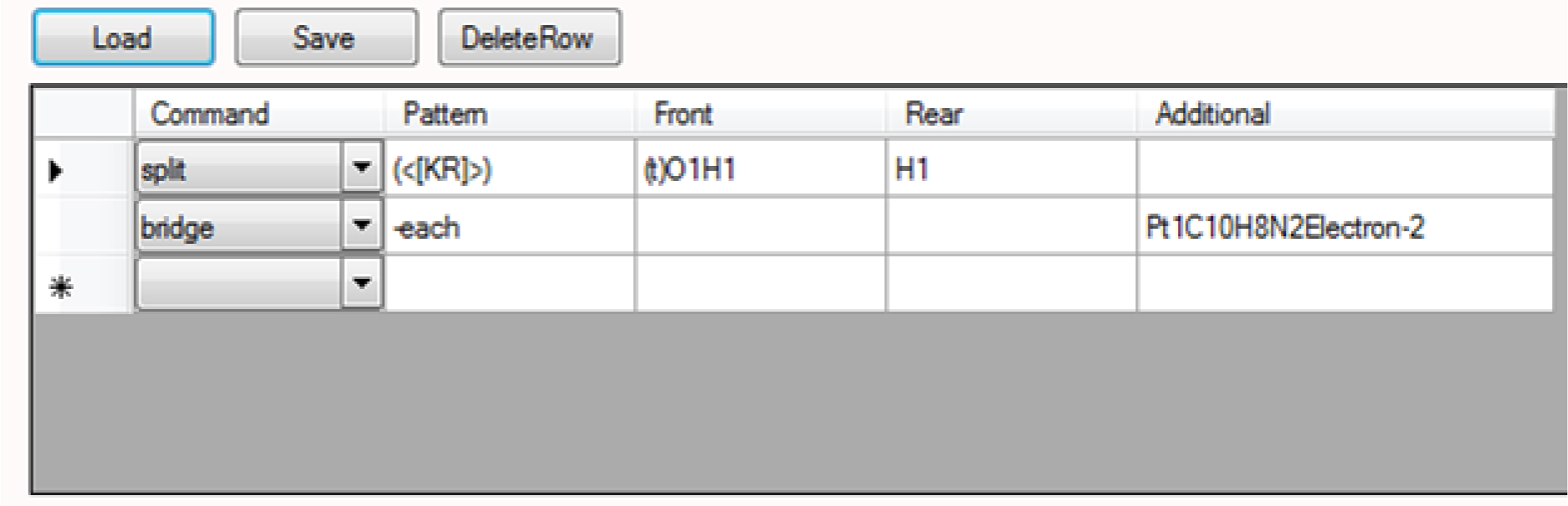
To generate a list of theoretical peptides from complete hydrolysis of the entered protein sequence (figure 3.7) at peptide bonds on the C-terminal side of Lys <u>and</u> Arg residues. The list contains additional chemical species consisting of a generated peptide possessing a Pt(bpy)^2+^ centre or any two of the generated peptides bridged by a Pt(bpy)^2+^ centre.

**Figure A2.15.**
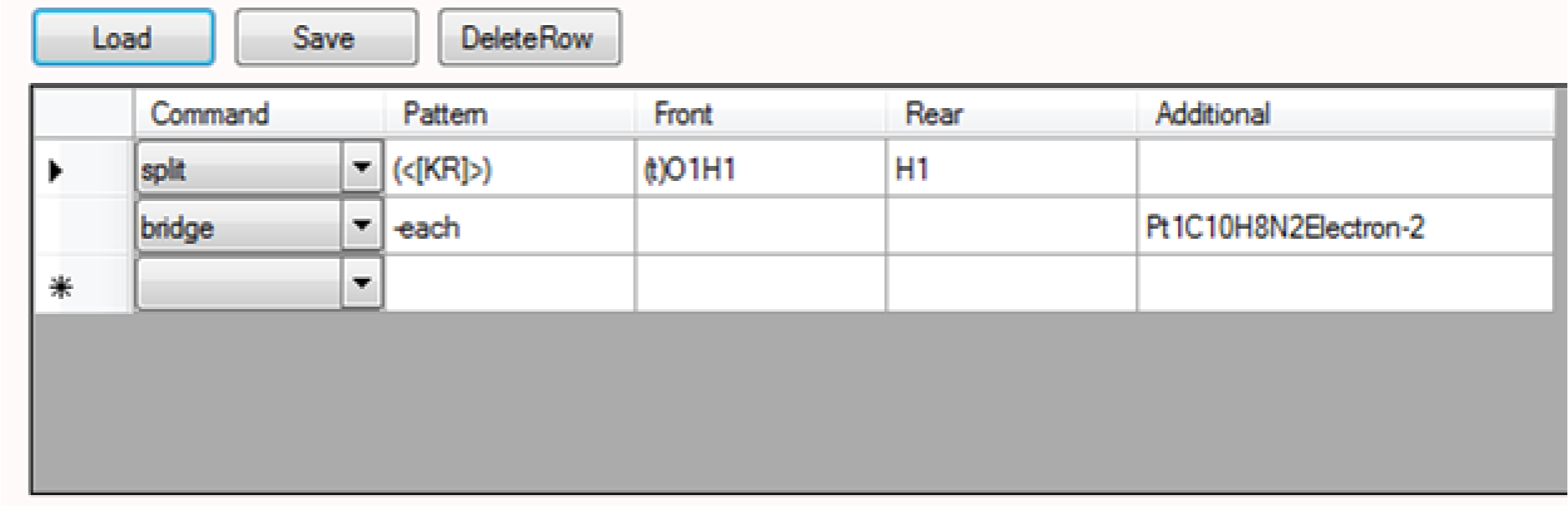
To generate a list of theoretical peptides from complete hydrolysis of the entered protein sequence (figure 3.7) at peptide bonds on the C-terminal side of Lys <u>and</u> Arg residues. The list contains additional chemical species consisting of a generated peptide possessing a Pt(bpy)2+ centre or any two of the generated peptides bridged by a Pt(bpy)2+ centre, if the peptide(s) has a His or Met residue.

